# Emotional tagging retroactively promotes memory integration through rapid neural reactivation and reorganization

**DOI:** 10.1101/2020.09.09.285890

**Authors:** Yannan Zhu, Lingke Zhang, Yimeng Zeng, Changming Chen, Guillén Fernández, Shaozheng Qin

## Abstract

Neutral events preceding emotional experiences are thought to be better remembered by tagging them as significant to simulate future event predictions. Yet, the neurobiological mechanisms how emotion transforms initially mundane events into strong memories remain unclear. By two behavioral and one fMRI studies with adapted sensory preconditioning paradigm, we show rapid neural reactivation and reorganization underlying emotion-tagged retroactive memory enhancement. Behaviorally, emotional tagging enhanced initial memory for neutral associations across the three studies. Neurally, emotional tagging potentiated reactivation of overlapping neural traces in the hippocampus and stimulus-relevant neocortex. Moreover, it induced large-scale hippocampal-neocortical reorganization supporting such retroactive benefit, as characterized by enhanced hippocampal-neocortical coupling modulated by the amygdala during online processing, and a shift from stimulus-relevant neocortex to transmodal prefrontal-parietal areas during offline post-tagging rest. Together, emotional tagging retroactively promotes associations between past neutral events through stimulating rapid reactivation of overlapping representations and reorganizing related memories into an integrated network.

## Introduction

Emotion shapes learning and memory for our episodic experiences. Experiencing an emotional event such as a psychological trauma, for instance, often not only strengthens our memory for the event itself, but also can transform other mundane events that occur close in time into strong memories (K. S. LaBar & Cabeza, 2006; Tambini, Rimmele, Phelps, & Davachi, 2017). There has been substantial progress in understanding the mechanisms underlying memory enhancement for emotional events themselves, through autonomic reactions to emotional arousal stimulating neural ensemble representations encoding what refers to emotional memory (Hamann, 2001; K. S. LaBar & Cabeza, 2006). However, our understanding of the mechanisms how emotional arousal reorganizes memories in a way that transforms initially mundane information into strong memories is still in its infancy. In many circumstances, the significance of our experiences such as reward or punishment, often occurs after the event. Since we cannot determine which event will be important later, human episodic memory is theorized to organize our experiences into a highly adaptive network of malleable representations, that can be prioritized in terms of the significance of preceding or following events (Dunsmoor, Murty, Davachi, & Phelps, 2015; Ritchey, Murty, & Dunsmoor, 2016). Such a mechanism allows seemingly mundane events to take on significance following a new salient experience, thereby generalizes their memories to future use in ever-changing environment. However, this retroactive effect may also lead to maladaptive generalization recognized as a cognitive hallmark of core symptoms in some mental disorders, like posttraumatic stress disorder (PTSD) and phobia (Lange et al., 2019; Mary et al., 2020). Deciphering the neurobiological mechanisms of emotion-induced retroactive memory enhancement in humans is thus critical for further understanding of maladaptive generalization in these disorders.

In past decades, rodent work has provided compelling evidence for retroactive memory enhancement following a significantly salient or emotional experience known as behavioral or emotional tagging (Takeuchi et al., 2016; Wang, Redondo, & Morris, 2010). At the molecular and cellular levels, an influential synaptic tagging and capture model (STC) proposes a mechanism, by which initially weak memories are strengthened by subsequent strong activation engaging overlapping neural ensembles with long-term potentiation (LTP) processes minutes to a few hours later (Frey & Morris, 1997; Redondo & Morris, 2011). Animal models typically opt for a state-like learning paradigm involving novelty exploration as strong input, which leads to an overall retroactive enhancement of weak memories encoded within a certain time window (Redondo & Morris, 2011; Takeuchi et al., 2016). In our everyday memory, however, the emotional effects often occur on some specific episodic memory representing a selectively single-shot learning. It thus remains an open question whether the conventional STC model based on state-like tagging paradigm can readily account for the effects of trial-specific learning or tagging on human episodic memory. At a systems level, evidence from animal and human studies brings memory allocation and integration models together to suggest that a shared network of neural ensembles links distinct memories encoded close in time or related in meaning (Margaret L. Schlichting & Frankland, 2017). By this view, there appears an integrative encoding mechanism by which a newly encoded event can be updated and reorganized into relevant episodic memory through reactivation of overlapping neural ensembles engaged in both initial and new learning (M. L. Schlichting & Preston, 2014; Shohamy & Wagner, 2008). Thus, it is possible that a retroactive memory benefit emerges rapidly for specific neutral information that occurs before emotional tagging. However, the neurobiological mechanisms how such trial-specific emotional tagging rapidly reorganizes human episodic memory remain elusive.

Memory allocation and integration models posit that related events are integrated into episodic memory through co-activation of overlapping neural ensembles (Margaret L. Schlichting & Frankland, 2017; Silva, Zhou, Rogerson, Shobe, & Balaji, 2009). Human neuroimaging studies provide compelling evidence supporting the integrative encoding mechanism, by which memories are organized into an integrated memory network across associations with overlapping representations (M. L. Schlichting & Preston, 2015; Shohamy & Wagner, 2008; van Kesteren, Brown, & Wagner, 2016). The hippocampus serves as an integrative hub by binding disparate representations in stimulus-selective neocortical areas into episodic memories (Kuhl, Shah, DuBrow, & Wagner, 2010; van Kesteren et al., 2016). Reactivation of hippocampal representations, together with hippocampal– neocortical coordinated interactions, has been well proposed to promote systems-level memory integration (M. L. Schlichting & Preston, 2014; Sutherland & McNaughton, 2000). The reactivation of overlapping neural traces is considered as a scaffold for integrating newly learnt information into existing memories, making it possible to reshape memories according to their future significance. However, it remains unknown how such reactivation contributes to the transformation of initially weak memories into strong memories following emotional tagging.

Previous studies measure memory reactivation using neural activation on overlapping brain regions involved in both old and new learning (Kuhl et al., 2010; Wimmer & Shohamy, 2012). This approach has provided useful information to identify specific brain systems and their activities during reactivation, but it offers limited insights into how distributed neural representations are reorganized following emotional learning. Multivariate pattern analysis of functional magnetic resonance imaging (fMRI) data has been used to assess multidimensional neural representations reflecting patterns of activation across neurons in specific brain areas during learning and memory (Norman, Polyn, Detre, & Haxby, 2006; Tambini & Davachi, 2019). Hence, we used multi-voxel pattern similarity to investigate how reactivation of shared neural representations with initial learning contributes to the retroactive benefit of emotional tagging on memory for neutral associations. Based on above neurobiological models and empirical observations in neuroimaging studies, we hypothesized that such emotion-tagged retroactive benefit would result from increased reactivation of hippocampal and stimulus-sensitive neocortical representations, which enhances the memory association between initial-learnt events and promotes their integration.

The amygdala, hippocampus and related neural circuits are known to play critical roles in memory enhancement for emotional experience (Dolcos, LaBar, & Cabeza, 2004; Richter-Levin, 2004). This emotion-induced memory enhancement is most likely based on autonomic reactions associated with emotional arousal, accompanying with (non)adrenergic signaling that modulates neural ensembles in the amygdala as well as the hippocampus and related regions involved in encoding emotional memories (Hamann, 2001; K. S. LaBar & Cabeza, 2006). Many studies, for instance, have suggested that emotional arousal can lead to better episodic memory through strengthening hippocampal-amygdala coupling during encoding and promoting hippocampal-neocortical dialogue for reactivation (de Voogd, Fernández, & Hermans, 2016; Richardson, Strange, & Dolan, 2004). Beyond its modulation on memory at the time of online encoding, emotional arousal is also thought to strengthen episodic memory by stimulating offline consolidation process during wakeful rest and sleep, involving reconfiguration of hippocampal functional connectivity with distributed neocortical networks (Kevin S LaBar & Phelps, 1998; McGaugh, 2018). Indeed, neuroimaging studies have demonstrated notable changes in hippocampal-neocortical connectivity during post-learning rest predictive of subsequent memory performance, confirming a possible role of early offline consolidation (Daniel L. Schacter, Benoit, & Szpunar, 2017; Margaret L Schlichting & Preston, 2016). However, it remains unknown how emotion-tagged online and offline mechanisms support retroactive benefits on episodic memory. Given the modulatory effect of emotion on memory at both encoding and consolidation, we further hypothesized that emotional tagging would retroactively promote memory integration for neutral events, by acting on not only hippocampal dialogue with amygdala and related neocortical regions during encoding, but also offline hippocampal-neocortical reorganization during post-tagging rest.

Here we tested these hypotheses by two behavioral studies and one event-related functional magnetic resonance imaging (fMRI) study using an adapted sensory preconditioning paradigm (Brogden, 1939). This paradigm distinguishes among the process involved in memory formation, the process involved in generalization of value information and process involved in plasticity underlying learning, allowing us to investigate mechanisms of different memory processes more precisely (Dunsmoor, White, & LaBar, 2011; Indovina, Robbins, Núñez-Elizalde, Dunn, & Bishop, 2011; Wimmer & Shohamy, 2012). In each of the three studies, the experimental design consisted of three phases: an initial learning, a following emotional tagging and a surprise associative memory test **(Fig. 1A)**. In Study 1 (N = 30), participants learnt 72 neutral face-object pairs during the initial learning phase. Thereafter, each face from the initial learning phase was presented as a cue and tagged by either an aversive screaming voice (aversive condition) or a neutral voice (neutral condition) during the emotional tagging phase. A surprise associative memory test was administered 30 minutes later to assess memory performance for initial face-object associations. In Study 2 (N = 28), the reproducibility of Study 1 was examined by another independent research group. In Study 3 (N = 31), participants underwent fMRI scanning during the initial learning and emotional tagging phases that were interleaved by three rest scans, with concurrent skin conductance recording to monitor autonomic arousal.

**Figure 1.**
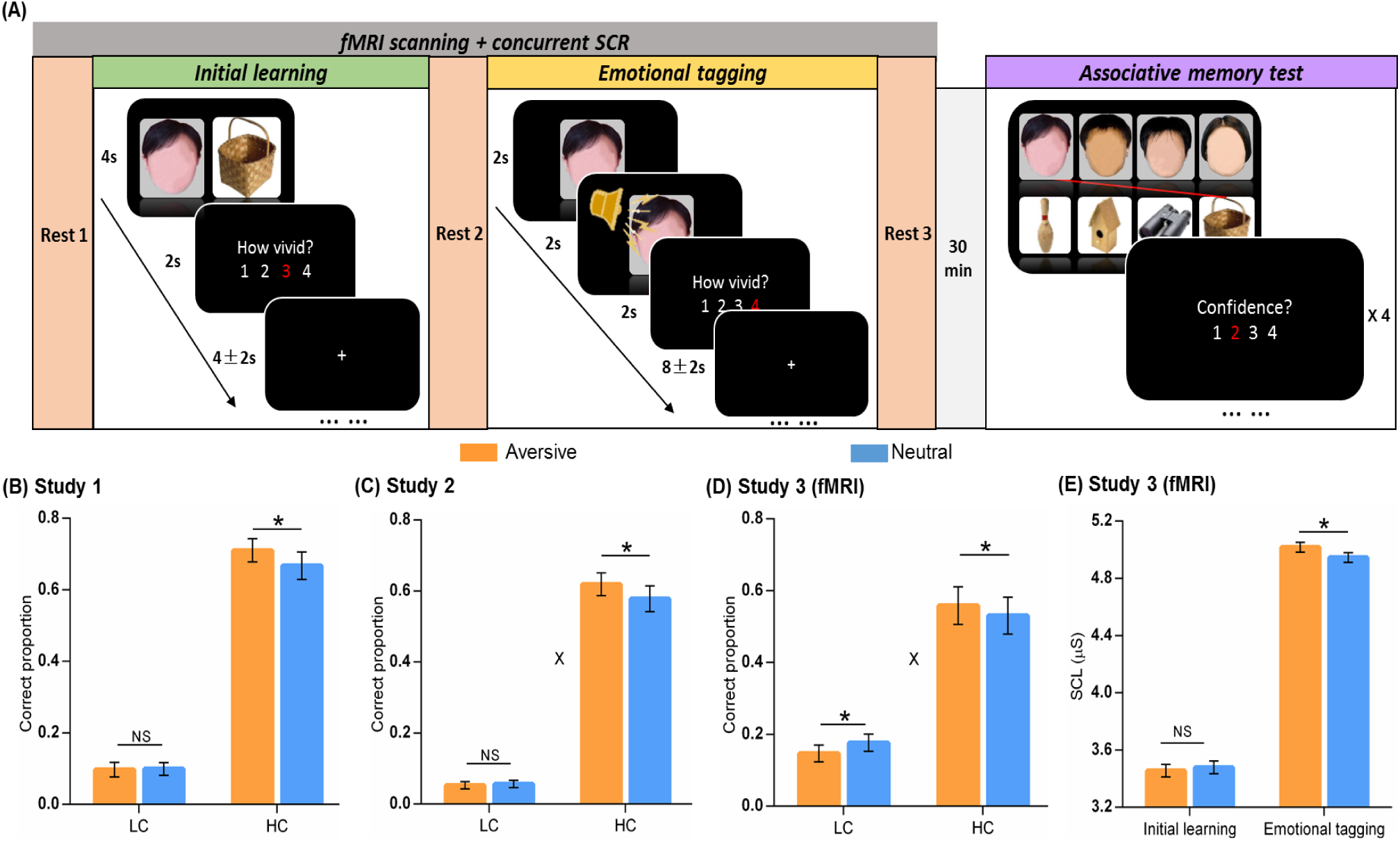
Experimental design and behavioral performance. **(A)** The experiment consisted of three phases. During initial learning phase, participants were instructed to vividly imagine each face and its paired object interacting with each other. During emotional tagging phase, each face of initial learning was presented again and then tagged with either an aversive scream or a neutral voice. Participants were also instructed to imagine each face and its tagged voice interacting. After a 30-min delay, participants performed a recognition memory test for face-object associations by matching each face (*top*) with one of the four objects (*bottom*). Faces in the figure are completely obscured for copyright reasons. **(B, C, D)** Associative memory performances in Study 1, 2 and 3 (fMRI). Bar graphs depict averaged correctness for face-object associations remembered with low (LC) and high (HC) confidence in aversive and neutral conditions. **(E)** Skin conductance levels (SCLs) in Study 3 (fMRI). Bar graphs depict averaged SCLs in aversive and neutral conditions during initial learning and emotional tagging phases separately. Error bars represent standard error of mean. “X” indicates significant interaction (*p* < 0.05). Notes: SCR, skin conductance recording; NS, non-significant; \**p* < 0.05; two-tailed tests.

A set of multi-voxel pattern similarity, and task-dependent/ task-free functional connectivity analyses, in conjunction with machine-learning prediction and structural equation modeling, were implemented to assess reactivation of initial learning activity, functional coupling during online emotional tagging and offline rests, and their relationships with emotion-induced retroactive memory enhancement. Consistent with our hypotheses, we found that emotional tagging retroactively enhanced memories for neutral associations initially learnt. This rapid retroactive enhancement was associated with prominent increases in reactivation of initial learning activity in the hippocampus and stimulus-sensitive neocortex, as well as strengthening hippocampal coupling with amygdala and related neocortical areas during emotional tagging and hippocampal-neocortical offline reorganization during post-tagging rest. Our findings suggest the neurobiological mechanisms by which memories for mundane neutral information can be prioritized when subsequently tagged with emotionally arousing experiences, through rapid reactivation of overlapping neural traces and reorganization of related memories into an integrated memory network according to their updated value weights.

## Results

### Emotional tagging retroactively enhances associative memory for initial neutral events

First, we examined the emotion-tagged retroactive effect on associative memory performance from two behavioral studies and one event-related fMRI study separately. One-sample t-tests on each confidence level revealed that the proportion of correctly remembered face-object associations with a 3 or 4 confidence rating (on a four-point rating scale: 4 = “Very confident”, 1 = “Not confident at all”) was significantly higher than chance across three studies (all *p* < 0.05; *Fig. S2 including statistics*). However, the proportion of remembered associations with a 1 or 2 rating was not reliably higher than chance (*Fig. S2 including statistics*). This indicates that remembering with high confidence may reflect more reliable and stronger memories as compared to low confident remembrance. Thus, all remembered associations were then sorted into a high confident bin with 3 and 4 ratings, and a low-confident bin with 1 and 2 ratings likely reflecting guessing or familiarity.

Separate 2 (Emotion: aversive vs. neutral) by 2 (Confidence: high vs. low) repeated-measures ANOVAs were conducted for three studies. For Study 1, this analysis revealed main effects of Emotion (F_(1,29)_ = 5.18, *p* = 0.030, partial η^2^ = 0.15) and Confidence (F_(1,29)_ = 142.07, *p* < 0.001, partial η^2^ = 0.83; **Fig. 1B**). Post-hoc comparisons revealed better memory for face-object associations only with high confidence (t_(29)_ = 2.18, *p* = 0.037, d_av_ = 0.22) in the aversive than the neutral condition. These results were reproducible by an independent research group in Study 2. Parallel analysis revealed main effects of Emotion (F_(1,27)_ = 5.53, *p* = 0.026, partial η^2^ = 0.17) and Confidence (F_(1,27)_ = 206.14, *p* < 0.001, partial η^2^ = 0.88), and an Emotion-by-Confidence interaction (F_(1,27)_ = 4.94, *p* = 0.035, partial η^2^ = 0.16; **Fig. 1C**). Post-hoc comparisons also revealed better associative memory with high confidence in the aversive (vs. neutral) than condition (t_(27)_ = 2.47, *p* = 0.020, d_av_ = 0.23). In Study 3 (fMRI), we again observed a similar pattern of results, including the main effect of Confidence (F_(1,27)_ = 29.35, *p* < 0.001, partial η^2^ = 0.52) and the Emotion-by-Confidence interaction (F_(1,27)_ = 8.61, *p* = 0.007, partial η^2^ = 0.24), especially with a retroactive enhancement on associative memory with high confidence in aversive (vs. neutral) condition (t_(27)_ = 2.59, *p* = 0.015, d_av_ = 0.10; **Fig. 1D**). Together, these results indicate an emotion-induced retroactive enhancement which only occurs on correctly remembered associations with high confidence in aversive condition.

To verify the elevation of autonomic arousal during emotional tagging and its potential effects on memory, we conducted separate paired-t tests for skin conductance levels (SCLs) in the fMRI study (Study 3). These analyses revealed higher SCL in the aversive than the neutral condition during emotional tagging phase (t_(27)_ = 2.10, *p* = 0.048, d_av_ = 0.22), but no evidence for a reliable difference during initial learning phase (t_(27)_ = −0.52, *p* = 0.610; **Fig. 1E**). These results indicate a higher level of automatic arousal induced by emotional tagging in the aversive than neutral condition, which is well likely to account for the emotion-induced retroactive memory enhancement.

### Emotional tagging potentiates reactivation of initial learning activity in the hippocampus and neocortex

Next, we investigated how emotional tagging affects reactivation of initial learning activity for face-object associations cued by overlapping faces (Study 3). To assess reactivation of neural activity, we estimated the similarity of stimulus-evoked multi-voxel activation patterns between the initial learning phase and the subsequent emotional tagging phase in aversive and neutral conditions separately (**Fig. 2A**). These analyses revealed significantly greater reactivation in the aversive than neutral condition in the bilateral hippocampus (t_(27)_ = 2.53, *p* = 0.017, d_av_ = 0.24; **Fig. 2B** *left*) and face/object-sensitive neocortical regions including the fusiform face area (FFA) and the lateral occipital cortex (LOC) (t_(27)_ = 2.99, *p* = 0.006, d_av_ = 0.17; **Fig. 2B** *middle*). In addition, we also implemented the whole-brain exploratory analysis with searchlight algorithm and identified other brain regions showing greater neural reactivation in the aversive than neutral condition (*Table S2*).

**Figure 2.**
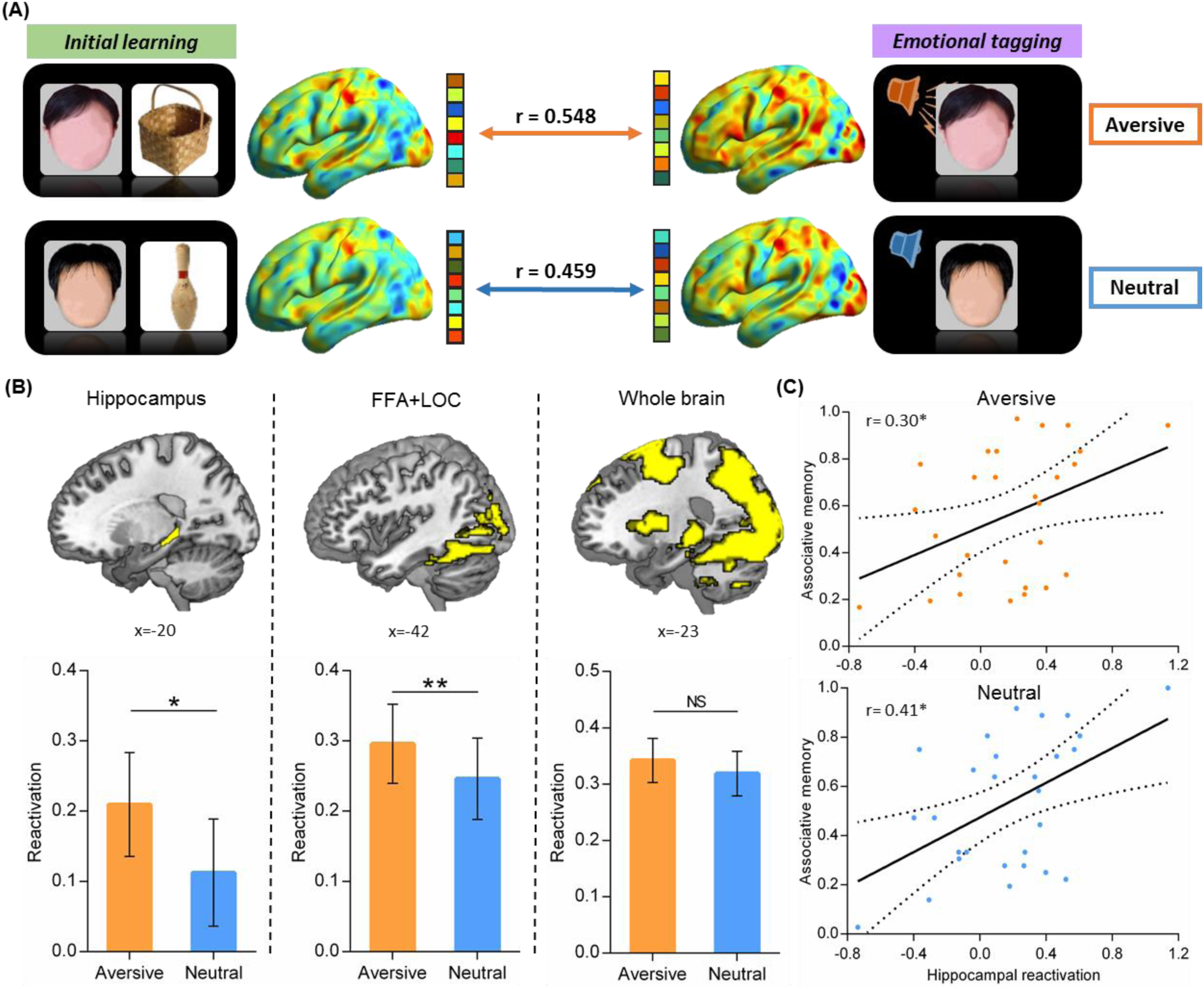
Neural reactivation of initial learning activity during emotional tagging. **(A)** An illustration of reactivation analysis by computing similarity of stimulus-evoked multi-voxel activity patterns between initial learning and emotional tagging phases. Example data from one subject is shown, with sagittal views of activation maps for aversive and neutral conditions during initial learning (*left*) and emotional tagging (*right*) phases. Faces in the figure are completely obscured for copyright reasons. **(B)** Bar graphs depict averaged reactivation in each condition on the hippocampus (*left*), face/object-sensitive neocortical regions (*middle*) including FFA and LOC, and a whole-brain activation mask (*right*). Error bars represent standard error of mean. **(C)** Scatter plots depict positive correlations of hippocampal reactivation with associative memory in aversive (*upper*) and neutral (*lower*) conditions. Dashed lines indicate 95% confidence intervals, and solid lines indicate the best linear fit. Notes: NS, not significant; \**p* < 0.05; \*\**p* < 0.01; FFA, face fusiform area; LOC, lateral occipital cortex.

Our further analyses revealed that greater neural reactivation occurred only after the onset of aversive voices, but not at the onset of face cues (*Fig. S3*). Interestingly, we observed no reliable difference of reactivation between two conditions in the whole-brain activation mask (t_(27)_ = 1.70, *p* = 0.10; **Fig. 2B** *right*), indicating that such emotion-induced increase in reactivation is not a global state across the whole brain. Critically, prediction analyses based on machine learning algorithms revealed that hippocampal reactivation was positively predictive of memory performance for face-object associations in both aversive (r_(predicted, observed)_ = 0.30, *p* = 0.041; **Fig. 2C** *upper*) and neutral conditions (r_(predicted, observed)_ = 0.41, *p* = 0.017; **Fig. 2C** *lower*). Together, these results indicate that emotional tagging potentiates reactivation of initial learning activity in the task-related hippocampus and stimulus-sensitive neocortex, with higher hippocampal reactivation predictive of better associative memory for initial neutral events in general.

### Emotional tagging enhances hippocampal coupling with the amygdala and neocortex

To further investigate how emotional tagging modulates the functional interactions of the hippocampus and its related neural circuits relevant for emotional episodic memory, we conducted task-dependent psychophysiological interaction (PPI) analyses for initial learning and emotional tagging phases separately to assess functional connectivity of the hippocampal seed with every other voxel of the brain (**Fig. 3A**). We mainly focused on hippocampal functional connectivity with the amygdala (**Fig. 3B**) and face/object-sensitive neocortical regions (i.e., FFA and LOC; **Fig. 3C**).

**Figure 3.**
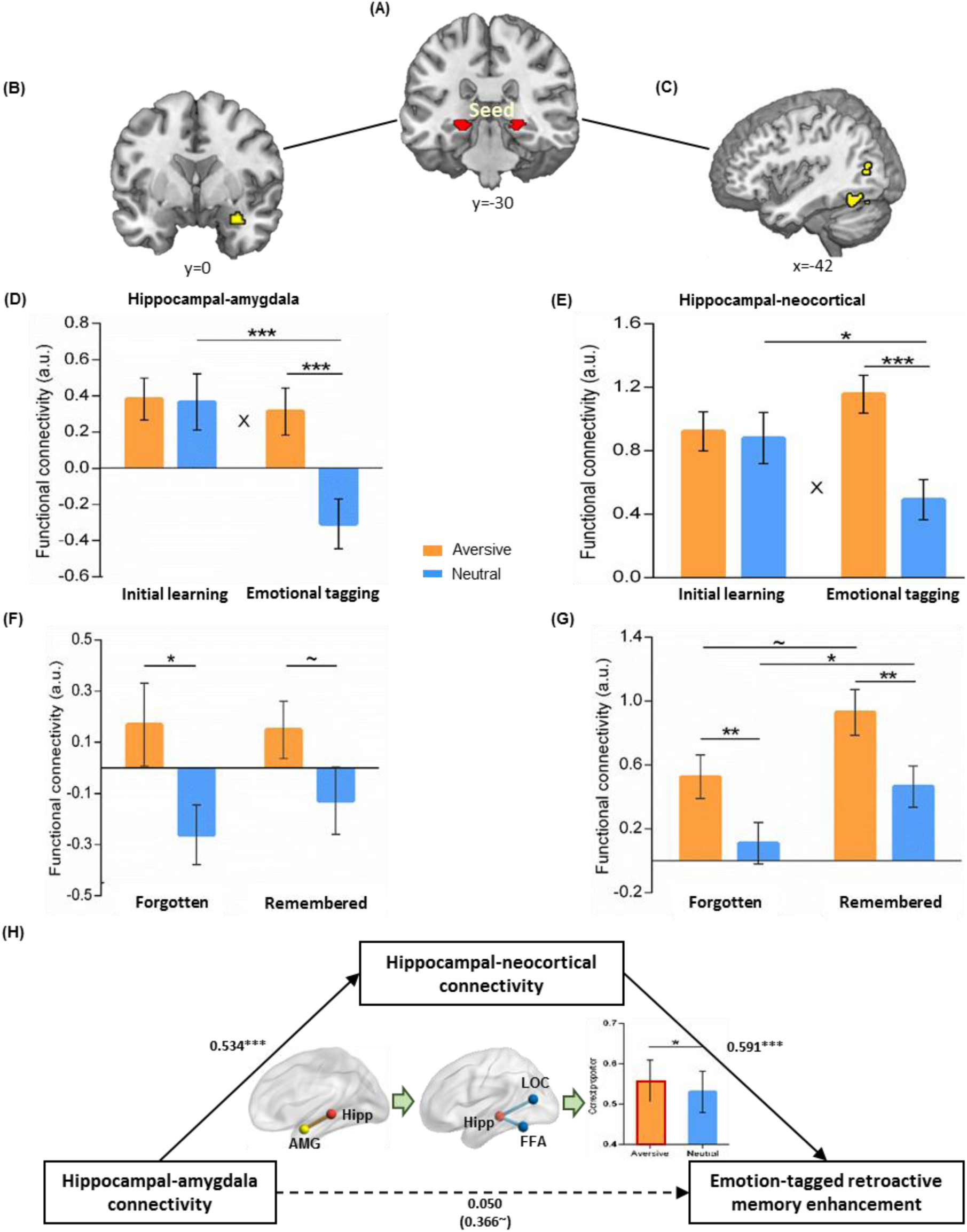
Online task-dependent hippocampal connectivity with the amygdala and neocortical regions in relation to emotion-tagged retroactive memory enhancement. **(A)** The bilateral hippocampal seed used in task-dependent gPPI functional connectivity analyses. **(B)** Significant cluster in the right amygdala, and **(C)** significant clusters in the left face/object-sensitive neocortical regions, showing greater functional coupling with the hippocampus in aversive (versus neutral) condition during emotional tagging. **(D, E)** Bar graphs depict averaged hippocampal connectivity with the amygdala and neocortical regions in aversive and neutral conditions during initial learning and emotional tagging separately. “X” indicates significant interaction. **(F, G)** Bar graphs depict averaged hippocampal connectivity with the amygdala and neocortical regions for face-object associations remembered with high confidence and forgotten in aversive and neutral conditions separately. Error bars represent standard error of mean. **(H)** The mediating effect of hippocampal-neocortical connectivity on the positive association between hippocampal-amygdala connectivity and emotion-tagged retroactive memory enhancement. Paths are marked with standardized coefficients. Solid lines indicate significant paths, and dashed lines indicate non-significant paths. The coefficient in bracket shows the correlation before hippocampal-neocortical connectivity was included into this model. Notes: ∼ *p* < 0.08; \**p* < 0.05; \*\**p* < 0.01; \*\*\**p* < 0.001; two-tailed tests; Hipp, hippocampus; AMG, amygdala.

Data extracted from these regions were submitted to separate 2 (Emotion: aversive vs. neutral) by 2 (Phase: initial learning vs. emotional tagging) repeated-measures ANOVAs. These analyses revealed significant Emotion-by-Phase interaction effects in the amygdala and neocortical regions including FFA and LOC (both F_(1,27)_ > 8.17, p < 0.008, partial η^2^ > 0.23; **Fig. 3D-E**). Post-hoc comparisons revealed significantly greater hippocampal connectivity with the amygdala and neocortical regions in aversive than neutral condition during emotional tagging phase (both t_(27)_ > 5.20, *p* < 0.001, d_av_ > 0.85), but not during the initial learning phase (both t_(27)_ < 0.09, *p* > 0.799). Interestingly, there were significant decreases in hippocampal connectivity with the amygdala and neocortical regions from the initial learning phase to the emotional tagging phase in the neutral condition (both t_(27)_ < −2.52, *p* < 0.018, d_av_ > 0.50), but not in the aversive condition (both t_(27)_ > −0.39, *p* > 0.180). In addition, parallel analyses revealed a very similar pattern of results for hippocampal connectivity with the face-sensitive FFA and the object-sensitive LOC separately (*Fig. S4*). Hippocampal connectivity with these regions during emotional tagging phase was also measured as the change relative to baseline initial learning phase, indicating the relative hippocampal connectivity was still significantly higher in the aversive than neutral condition (all t_(27)_ > 2.67, *p* < 0.015, d_av_ > 0.63; *Fig. S5*). This pattern of results indicates that emotional tagging induces online task-dependent hippocampal functional reorganization with prominent increases in hippocampal-amygdala and hippocampal-neocortical couplings.

### Emotion-charged hippocampal coupling predicts associative memory for initial neutral events

We then investigated how emotion-charged hippocampal connectivity contributes to the retroactive benefits on associative memory. Data extracted from hippocampal connectivity with the amygdala and neocortical regions during the emotional tagging phase were submitted to separate 2 (Emotion: aversive vs. neutral) by 2 (Memory: remembered with high confidence vs. forgotten) repeated-measures ANOVAs. The analysis for hippocampal-amygdala connectivity revealed a reliable main effect of Emotion (F_(1, 26)_ = 20.11, *p* < 0.001, partial η^2^ = 0.44), but neither a main effect of Memory nor an Emotion-by-Memory interaction (both F_(1,26)_ < 0.30, *p* > 0.590; **Fig. 3F**). Post-hoc comparisons revealed that hippocampal-amygdala connectivity was greater in aversive than neutral condition, generally for face-object associations both later remembered with high confidence (t_(26)_ = 1.91, *p* = 0.068, d_av_ = 0.44) and forgotten (t_(26)_ = 2.47, *p* = 0.021, d_av_ = 0.59). Parallel analysis for hippocampal-neocortical connectivity revealed significant main effects of Emotion (F_(1,26)_ = 18.31, *p* < 0.001, partial η^2^ = 0.41) and Memory (F_(1, 26)_ = 5.39, *p* = 0.028, partial η^2^=0.17), but not an Emotion-by-Memory interaction (F_(1,26)_ = 0.06, *p* = 0.802; **Fig. 3G**). Post-hoc comparisons revealed that hippocampal-neocortical connectivity was greater in aversive than neutral condition (remembered: t_(26)_ = 2.85, *p* = 0.008, d_av_ = 0.66; forgotten: t_(26)_ = 3.53, *p* = 0.002, d_av_ = 0.60), and for face-object associations remembered with high confidence than forgotten (aversive: t_(26)_ = 1.90, *p* = 0.069, d_av_ = 0.55; neutral: t_(26)_ = 2.15, *p* = 0.041, d_av_ = 0.53).

Critically, we found that greater hippocampal connectivity with significant clusters in the amygdala and neocortical regions (remembered with high confidence relative to forgotten) were predictive of better associative memory in aversive condition (amygdala: r_(predicted, observed)_ = 0.24, *p* = 0.076; neocortical regions: r_(predicted, observed)_ = 0.59, *p* = 0.002), but not in neutral condition (both r_(predicted, observed)_ < 0.11, *p* > 0.180). In addition, parallel analyses were conducted for hippocampal connectivity with significant clusters in the FFA and LOC separately, which revealed very similar result patterns from both ANOVA and prediction analyses (*Fig. S6*). These results indicate that emotion-charged hippocampal functional coupling with the amygdala and neocortical regions positively predicts associative memory for initial neutral events.

### Emotion-tagged retroactive memory enhancement via task-dependent functional coupling between the hippocampus, amygdala and neocortex

We further investigated how functional organization of the hippocampus, amygdala and neocortical systems during emotional tagging contributes to the emotion-induced retroactive benefits on associative memory, by implementing structural equation modeling (SEM) to assess the mediating pathways of hippocampal-amygdala and hippocampal-neocortical connectivity with associative memory outcomes. We constructed a full mediation model with hippocampal functional connectivity (remembered with high confidence relative to forgotten) and individual’s associative memory performance (correctness proportion with high confidence) in the aversive condition (**Fig. 3H**). The path coefficient for hippocampal-amygdala connectivity to emotion-tagged associative memory dropped to non-significance when the mediating path was added. The mediating effect of hippocampal-neocortical connectivity on the association between hippocampal-amygdala connectivity and associative memory in aversive condition was significant (Indirect Est = 0.32, *p* = 0.004, 95% CI = [0.098, 0.533]). However, owing to non-significant relationship of hippocampal connectivity with associative memory in the neutral condition, neither path coefficients nor the mediating effect in the mediation model was significant (*Fig. S7*). In addition, parallel SEM analyses revealed that hippocampal-amygdala connectivity positively predicted emotion-induced increase in neural reactivation in hippocampus and neocortical regions, which was also mediated by hippocampal-neocortical functional connectivity (*Fig. S8*). These results indicate that hippocampal-amygdala connectivity positively predicts both emotion-induced retroactive enhancement on associative memory and increased reactivation in hippocampus and neocortical regions, which is mediated by hippocampal-neocortical functional coupling during emotional tagging.

### Emotion-tagged retroactive memory enhancement is associated with the reorganization of hippocampal connectivity during post-learning rest

Hippocampal-neocortical reorganization is implicated in systems consolidation (Liu et al., 2016; S. Qin, Cho, et al., 2014) and thus, we tested the hypothesis whether emotional tagging would modulate hippocampal connectivity during post-learning rest. We implemented seed-based correlational analyses of resting-state fMRI data separately for three rest scans that interleaved the initial learning and emotional tagging phases, with the hippocampus as seed to assess its functional connectivity with the rest of the brain (**Fig. 4A**). We computed difference maps of hippocampal connectivity during post-learning rest after initial learning (relative to before: Rest 2 vs. 1), as well as during post-tagging rest after emotional tagging (relative to before: Rest 3 vs. 2). Thereafter, we associated these connectivity maps with individual effects on memory performance (i.e., memory performance for neutral associations in the aversive relative to neutral condition). These analyses revealed a significant cluster in the lateral occipital cortex (LOC) after initial learning relative to baseline (**Fig. 4B**; *Table S3*), with greater hippocampal-LOC connectivity predictive of better emotion-tagged memory enhancement (r_(predicted, observed)_ = 0.53, *p* = 0.002; **Fig. 4D**). More importantly for the question at issue, when comparing rest after emotional tagging relative to before, we found significant clusters in a set of brain regions including the anterior prefrontal cortex (aPFC), the inferior parietal lobule (IPL) extending into the angular gyrus (AG), and the posterior cingulate cortex (PCC) (**Fig. 4C**; *Table S3*), with greater hippocampal connectivity with these regions predicting larger emotion-tagged memory benefit (aPFC: r_(predicted, observed)_ = 0.51; IPL: r_(predicted, observed)_ = 0.53; PCC: r_(predicted, observed)_ = 0.50; all *p* < 0.005; **Fig. 4E**). These results indicate a prominent shift of offline hippocampal connectivity from the lateral occipital cortex to more distributed transmodal prefrontal and posterior parietal areas, which associates with emotion-tagged retroactive memory enhancement.

**Figure 4.**
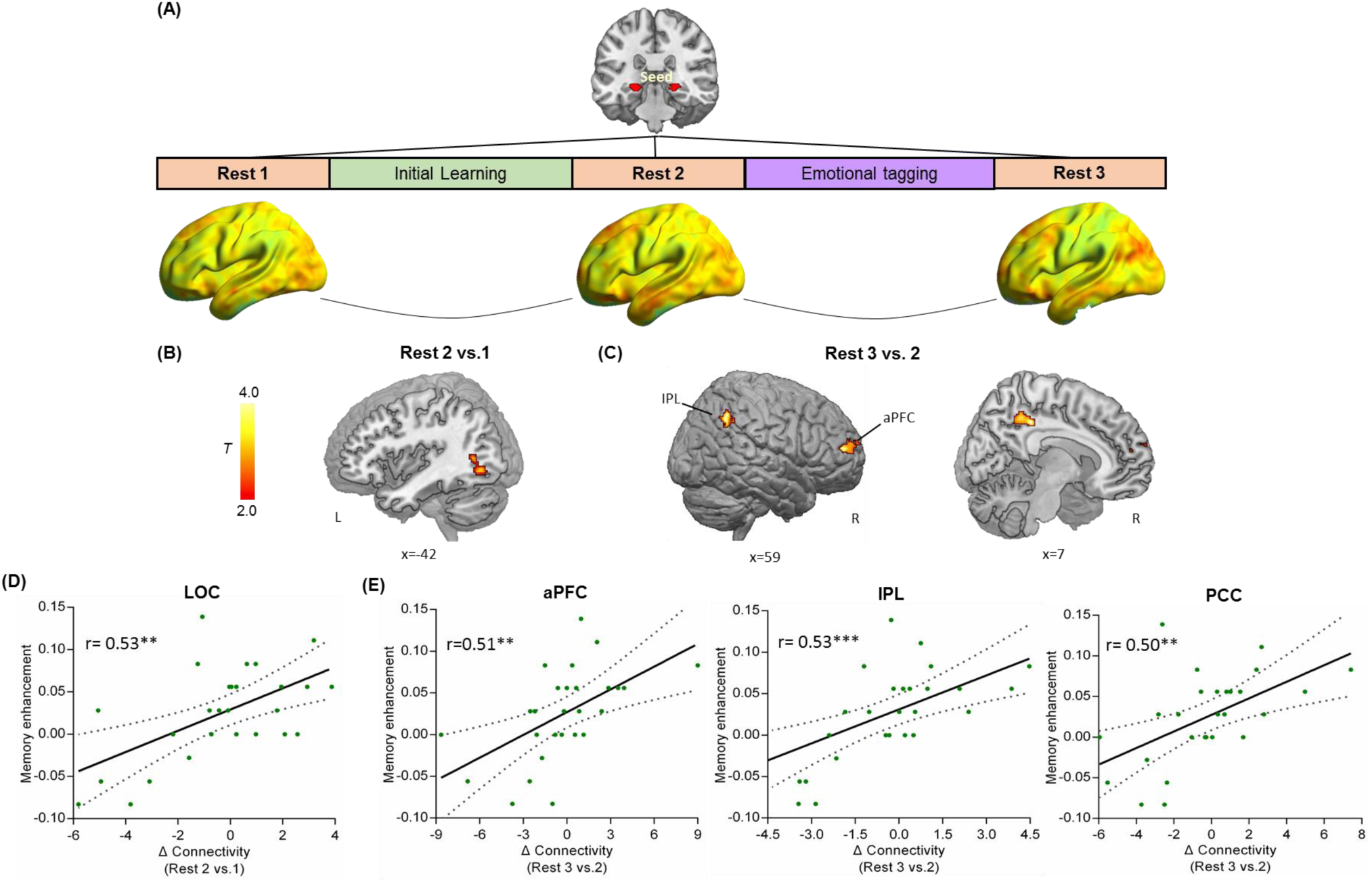
A prominent shift of hippocampal-cortical connectivity during post-learning rest in relation to emotion-tagged retroactive memory enhancement. **(A)** An illustration of hippocampal-seeded functional connectivity analyses, and sagittal views of hippocampal connectivity maps at the group level for three rest scans. **(B)** Significant cluster in the lateral occipital cortex (LOC), showing its greater connectivity with the hippocampus at Rest 2 relative to Rest 1. **(C)** Significant clusters in the right anterior prefrontal cortex (aPFC), the bilateral inferior parietal lobule (IPL) extending into angular gyrus and the right posterior cingulate cortex (PCC), showing their greater connectivity with the hippocampus at Rest 3 relative to Rest 2. **(D-E)** Scatter plots depict positive correlations between emotion-tagged retroactive memory enhancement and hippocampal connectivity with the LOC after initial learning (Rest 2 versus 1), as well as aPFC, IPL and PCC after emotional learning (Rest 3 versus 2). Dashed lines indicate 95% confidence intervals, and solid lines indicate the best linear fit. Notes: Color bars represent T values; \*\**p* < 0.005; \*\*\**p* < 0.001; two-tailed tests.

## Discussion

By three studies, we investigated the neurobiological mechanisms of how memories for associated mundane events are retroactively modulated by following learning of emotional events. As expected, emotional tagging retroactively and selectively enhanced associative memory for initial neutral events. This rapid retroactive enhancement was associated with increased reactivation of initial learning activity in the hippocampus and stimulus-sensitive neocortex, as well as strengthened hippocampal coupling with the amygdala and neocortical regions during emotional tagging. Critically, hippocampal-amygdala coupling positively predicted the retroactive memory benefit and increased neural reactivation, which were both mediated by prominent increases in hippocampal-neocortical interactions. Moreover, we found a prominent shift of hippocampal-neocortical connectivity during offline post-tagging rest that was associated with emotion-tagged retroactive memory enhancement, from local stimulus-sensitive neocortex to more distributed transmodal prefrontal and posterior parietal regions. Our findings suggest that emotional tagging retroactively promotes the integration for past neutral events into episodic memory to foster future event predictions, through stimulating reactivation of overlapping memory traces and reorganization of related memories with their value weights in the relevant neural circuits.

### Emotion-tagged retroactive enhancement on memory integration for initial neutral information

Behaviorally, we observed that emotional tagging retroactively enhanced associative memory for initial neutral events. The rapid enhancement was selective for face-object associations that were later tagged with screaming voice in the aversive condition, but not in the neutral (control) condition. Notably, this effect was reproducible across two independent studies indicating its reliability and robustness. It is in line with the classical sensory preconditioning effects reported in previous studies, that a salient event would spread its value to potentially but indirectly associated mundane event and retroactively prioritize its memory or guide a decision on it (Dunsmoor et al., 2011; Wimmer & Shohamy, 2012). Beyond these studies focusing on item memory of indirectly associated event, our results provide novel evidence that the value of emotional event would also be directly tagged to the initial associations of neutral events during reactivation of overlapping memory traces. Thus, the subsequent salient information may promote the associations between past neutral events and reorganize related memories into a more tightly integrated network.

It is worth noting that we adopted auditory stimuli as emotional tagging to condition neutral image associations. The emotional tagging mainly aims to deliver value information to related mundane events, but not replace those old experiences as alternatives. Thus, it should be distinguished from a phenomenon, that new emotional learning potentially impacts its overlapping old experiences and disturb the coherence of related memories, through transient increases in acetylcholine and thereafter hippocampal pattern separation (Bisby, Horner, Bush, & Burgess, 2018; Decker & Duncan, 2020; Kuhl et al., 2010).

Nevertheless, this rapid effect differs from the retroactive memory enhancement previously reported in several humans and rodent studies under the framework of synaptic tagging-and-capture (STC) model (Braun, Wimmer, & Shohamy, 2018; Dunsmoor et al., 2015; Moncada & Viola, 2007). Because the STC model emphasizes an essential role of systems consolidation typically after several hours and even days to yield the retroactive memory benefits at the behavioral level (Frankland & Bontempi, 2005; Redondo & Morris, 2011; Wang et al., 2010). Our behavioral finding offers new insights into such emotion-induced retroactive benefits, that the emotional tagging with autonomic arousal occurring exactly during reactivation of initial memory may stimulate a rapid memory reorganization during both online learning and offline rest. Moreover, the rapid retroactive benefit shown in our study was pertaining to newly acquired neutral associations, which extends previous findings focusing on concept-related events that already existed in long-term memory schema (Dunsmoor, Martin, & LaBar, 2012; Dunsmoor et al., 2015).

### Emotion-charged reactivation of overlapping neural traces in the hippocampus and neocortex

In parallel with above described rapid, retroactive memory enhancement, our imaging results showed transient increases in neural pattern similarity between initial learning and emotional tagging phases in the aversive (vs. neutral) condition. This similarity measure is thought to reflect the degree of reactivation of overlapping neural traces triggered by each face cue, according to previous studies assessing the similarity of neural activity between encoding and retrieval (Jonker, Dimsdale-Zucker, Ritchey, Clarke, & Ranganath, 2018; Tompary & Davachi, 2017). It is thus likely that our observed increases in neural pattern similarity after the onset of aversive (vs. neutral) voices during emotional tagging could be caused by emotion-induced autonomic arousal accompanied with elevated catecholamine release, which rapidly potentiates neuronal excitability in currently activated brain regions like the hippocampus and stimulus-sensitive neocortical regions (Rogerson et al., 2014; Zhou et al., 2009). In other words, there appears to be a cued reactivation of overlapping neural traces with concurrent autonomic reactions related to arousal triggered by the aversive voices during emotional tagging. Based on memory allocation and integration models in rodents (Margaret L. Schlichting & Frankland, 2017; Silva et al., 2009), co-activation of overlapping neurons may serve as a mechanism by which memories for prior and present experiences can be allocated into an integrated representation. By this view, autonomic arousal triggered by emotional tagging may stimulate the excitability of overlapping neuronal ensembles that could strengthen the associations between prior memories for neutral information in an integrated memory network.

Moreover, we found prominent effects in the hippocampus and specific neocortical regions including FFA and LOC. This finding implies that the emotion-induced increase in reactivation of overlapping neural traces mainly occurs in those regions critical for episodic memory of faces and objects (Sutherland & McNaughton, 2000; Zeithamova, Dominick, & Preston, 2012). The hippocampus, for instance, is known to play a critical role in the integration of distributed information into episodic memory through a pattern completion process (Eichenbaum, 2004; Staudigl & Hanslmayr, 2018). Thus, hippocampal reactivation may strengthen the coherence of related representations throughout our brain (Kuhl et al., 2010; Tambini & Davachi, 2019; Wimmer & Shohamy, 2012). Indeed, we observed that hippocampal reactivation was predictive of memory for initially learnt face-object associations. The emotion-charged reactivation was also observed in FFA and LOC sensitive to faces and objects respectively, which is thought to integrate the representations of these events into a distributed neocortical network (Frankland & Bontempi, 2005). Together, our findings suggest that emotion-induced transient increases in reactivation of overlapping neural traces in the hippocampus and related neocortical regions may promote a rapid memory reorganization of prior and present events.

### Emotion-induced memory reorganization via hippocampal-amygdala-neocortical interactions

Coinciding with transient increases in reactivation, our results further suggest an emotion-induced memory reorganization by stimulating coupling between hippocampus, amygdala, and neocortical circuits during emotional tagging, as well as a prominent shift of hippocampal-neocortical connectivity from local stimulus-sensitive occipital area to more distributed prefrontal and posterior parietal systems during post-tagging rest. Four aspects of our data support this interpretation. First, we observed emotion-induced increases in hippocampal functional connectivity with the amygdala and face/object-sensitive neocortical regions in the aversive compared to neutral condition during emotional tagging. Second, we observed that both hippocampal-amygdala connectivity and hippocampal–neocortical connectivity during emotional tagging positively predicted the memory performance for face-object associations in the aversive, but not the neutral condition. It is known that hippocampal functional coordination with relevant neocortical regions plays a critical role in reactivation and integration of episodic memories (Nadel, Samsonovich, Ryan, & Moscovitch, 2000; Sutherland & McNaughton, 2000). Thus, this finding suggests a possible mechanism of emotion-induced online memory reorganization via hippocampal–neocortical functional coupling, coinciding with the modulation of amygdala activity.

Third and more importantly, we found that the positive relationship between hippocampal-amygdala coupling and emotion-induced retroactive memory enhancement was fully mediated by an increase in hippocampal-neocortical coupling. This finding proved the modulatory role of the amygdala acting on hippocampal functional coordination with the neocortex (i.e., FFA and LOC). This modulatory role may support more efficient information transmission and communication between the hippocampus and related neocortical regions (Damasio, 1989), and thereby lead to rapid reactivation of overlapping neural traces with past experiences and reorganize their value weights in an integrated memory network according to the significance of subsequent related events. This interpretation was further supported by our observations that hippocampal-neocortical coupling also mediated the positive relationships of hippocampal-amygdala coupling with reactivation in hippocampal and neocortical regions. Last but not least, we further observed notable hippocampal-neocortical reorganization during post-tagging rests predictive of emotion-tagged retroactive memory enhancement, with a prominent shift away from local stimulus-sensitive lateral occipital cortex to more distributed prefrontal and posterior parietal regions including the aPFC, PCC and IPL expanding into the AG. These transmodal prefrontal-parietal regions are core nodes of the default mode network (DMN) that plays a crucial role in remembering past events and imagining or simulating possible future use (D. L. Schacter, Guerin, & St Jacques, 2011; Spreng & Schacter, 2012). Thus, our findings suggest that emotional tagging not only rapidly strengthened online task-dependent hippocampal-neocortical coupling likely through amygdala modulation, but it also potentiated offline hippocampal-neocortical reorganization integrating memory traces into more distributed networks and prioritize them for future use. Taken together, our findings point toward a rapid emotion-modulated reorganization mechanism, by which memories for neutral events can be updated according to the significance of subsequent events. This emotion-modulated reorganization not only contributes to the integration of past and current experiences into episodic memory, but also supports future simulations.

**In conclusion**, our study demonstrates that emotional tagging can retroactively promote memory integration for preceding neutral events through an emotion-charged rapid memory reorganization mechanism, characterized by transient increases in reactivation of overlapping neural traces, strengthened online hippocampal-neocortical functional coupling modulated by the amygdala, and a shift of hippocampal-neocortical offline connectivity from local stimulus-sensitive neocortex to more distributed prefrontal and posterior parietal areas during post-tagging rest. These findings advance not only our understanding of the neurobiological mechanisms by which emotion can reshape our episodic memory of past neutral events fostering its adaptive priority in the future, but it may also provide novel insights into maladaptive generalization in mental disorders like PTSD.

## Methods

### Participants

A total of 89 young, healthy college students participated in three separate studies. In Study 1, 30 participants (16 females; mean age ± s.d., 22.23 ± 2.05 years old, ranged from 18 to 26) were recruited from Beijing area to participate in a behavioral experiment. In Study 2 by an independent research staff, 28 participants (14 females; mean age ± s.d., 21.83 ± 1.93 years old, ranged from 18 to 26) were recruited from Xinyang city in Henan province for a replication experiment to ensure the reliability of our behavioral findings from Study 1. In Study 3, another independent cohort of 31 participants (17 females; mean age ± s.d., 22.55 ± 2.25 years old, ranged from 18 to 27) was recruited from Beijing area to participate in an event-related fMRI experiment. Data from three participants were excluded from further analyses due to either falling asleep during fMRI scanning (n = 2) or poor memory performance (n = 1). All participants were right-handed with normal hearing and normal or corrected-to-normal vision, reporting no history of any neurobiological diseases or psychiatric disorders. Informed written consent was obtained from each participant before the experiment. The Institutional Review Board for Human Subjects at Beijing Normal University, Xinyang Normal University and Peking University approved the procedures for Study 1, 2 and 3 respectively.

### Materials

One hundred and forty-four face images (72 males and 72 females) were carefully selected from a database with color photographs of Chinese individuals unknown to participants (Chen et al., 2012), under following criteria suggested by previous studies (Shaozheng Qin et al., 2007): direct gaze contact, no headdress, no glasses, no beard, etc. There was also no strong emotional facial expression in these faces, and no significant difference in terms of arousal, valence, attractiveness, and trustworthiness between male and female faces according to rating results in a previous study (Liu et al., 2016). Seventy-two object images were obtained from a website (http://www.lifeonwhite.com) or publicly available resources on the internet (Dunsmoor et al., 2015). All objects were common in life with neutral valence. Four short clips of female voices were carefully selected from an audio source website (www.smzy.com), with two aversive screams serving as emotional tagging manipulation and two neutral voices (“Ah” and “Eh”) as control. Acoustic characteristics of the four voice clips were measured and controlled using Praat (www.praat.org), including duration (2 seconds), frequency (in Hertz) and power (in decibel) (*Table S1*). Two aversive screams had significantly higher arousal and lower valence than neutral voices (all *p* < 0.001; *Fig. S1A*), as measured on a 9-point self-rating manikin scale (i.e., 9 = “Extremely pleasant” or “Extremely arousing”, 1 = “Not pleasant at all” or “Not arousing at all”). There was no significant difference in arousal or valence between two aversive (or neutral) voices (all *p* > 0.05).

Faces were randomly split into two sets of 72 images with half male and half female faces for each participant: one list was paired with objects to create 72 face-object pairs for the initial learning phase and the other was served as foils for face recognition memory test (face item memory performance is shown in *Fig. S1B*). Then, each face in face-object pairs was randomly paired with one of four voices to create 72 face-voice pairs and assigned into either an aversive condition (36 faces tagged with aversive screams) or a neural condition (36 faces tagged with neutral voices) during the emotional tagging phase.

### Experimental procedures

In each of the three studies, the experimental design consisted of three consecutive phases: an initial learning, a followed-up emotional tagging, and a surprise recognition memory test (**Fig. 1A**). During the initial learning phase, participants were instructed to view 72 face-object pairs in an incidental encoding task. During the emotional tagging phase, faces from the initial learning phase were presented again as cues and randomly tagged with either an aversive or a neutral voice. After a 30-minute delay, participants performed a surprise recognition memory test for face-object associations. In Study 3, participants underwent fMRI scanning with concurrent recording of skin conductance while they were performing initial learning and emotional tagging phases interleaved by three rest scans. *Specifically*, the fMRI experiment began with an 8-min baseline rest scan (Rest 1), followed by the initial learning and emotional tagging phases. Each of the two learning phases was followed by another 8-min resting state scan (Rest 2 and 3). During rest scans, participants were shown a black screen and instructed to keep awake with their eyes open. Then, the surprise associative memory test was performed outside the scanner.

### Initial learning task

During initial learning, 72 faces were randomly paired with 72 objects to create 72 face-object associations for each participant. Each association was centrally presented on the screen for 4 seconds, and followed by a vividness rating scale for 2 seconds. To ensure incidental memory encoding, participants were instructed to vividly imagine each face interacting with its paired object and give a vividness rating on their imagined scenario on a 4-point Likert scale (i.e., 4 = “Very vivid”, 1 = “Not vivid at all”). Trials were interleaved by a fixation with an inter-trial interval jittered from 2 to 6 seconds (i.e., 4 seconds on average with 2-second step). A total of 72 face-object pairs were viewed twice, which was split into two runs with 12 minutes each.

### Emotional tagging task

During emotional tagging, each face of initial learning was presented at the center of the screen for 2 seconds, and then followed by concurrent presentation of the same face tagged with either an aversive or a neutral voice for another 2 seconds. To ensure the consistency with initial incidental learning task, participants were again instructed to imagine the face interacting with its corresponding voice and give their vividness rating on a 4-point Likert scale for 2 seconds. After that, each trial was followed by a relatively long inter-trial interval jittered from 6 to 10 seconds (i.e., 8 seconds on average with 2-second step), to eliminate potential contamination of voice-induced emotional arousal among neighboring trials. Faces and voices were randomly paired across participants. Totally 72 face-voice pairs were presented only once in a pseudo-randomized order that no more than 2 voices from the same condition appeared in a row. The emotional tagging phase lasted 16.8 minutes.

### Surprise recognition memory test

After a 30-minute delay, participants were instructed to perform a self-paced recognition memory test for face-object associations. Each trial consisted of four faces from initial learning on the top of screen and their corresponding objects randomly located on the bottom of screen. A total of 72 encoded face-object pairs were randomly sorted in 18 slides with 4 associations each. Participants were asked to match each face with one of the four objects according to their remembrance of face-object associations, and then gave their confidence rating on a 4-point scale (i.e., 4 = “Very confident”, 1 = “Not confident at all”) for each choice. Participants were required to make choice for each face in an order from left to right, and carefully recall before they made the choice to avoid errors.

### Behavioral data analysis

Participants’ demographic data and behavioral performance on memory accuracy and confidence rating from surprise memory test were analyzed using Statistical Product and Service Solutions (SPSS, version 22.0, IBM). One sample t-tests were conducted to examine the reliable memory performance of face-object associations as compared to chance level for each confidence level (from 1 to 4). Then, all remembered associations were sorted into a high confident bin with levels 3 and 4, and a low-confident bin with levels 1 and 2 (Squire, Wixted, & Clark, 2007). Separate 2 (Emotion: aversive vs. neutral) by 2 (Confidence: high vs. low) repeated-measures ANOVAs were conducted in three studies to examine the emotion-tagging effects on performance which reflects reliable and strong memory but not guessing or familiarity.

### Skin conductance recording and analysis

Skin conductance was collected to assess autonomic arousal induced by aversive screaming (versus neutral) voices during emotional tagging. It was recorded simultaneously with fMRI scanning using a Biopac MP 150 System (Biopac, Inc., Goleta, CA). Two Ag/AgCl electrodes filled with isotonic electrolyte medium were attached to the center phalanges of the index and middle fingers of each participant’s left hand. The gain set to 5, the low-pass filter set to 1.0 Hz, and the high-pass filters set to DC (Indovina et al., 2011). Data were acquired at 1000 samples per second and transformed into microsiemens (μS) before further analyses. Given the temporal course of skin conductance in response to certain event, mean skin conductance levels (SCLs) were calculated for a period of 6 seconds after each stimulus onset.

### Imaging acquisition

Whole-brain imaging data were collected on a 3T Siemens Prisma MRI system in Peking University. Functional images were collected using a multi-band echo-planar imaging (mb-EPI) sequence (slices, 64; slice thickness, 2 mm; TR, 2000 ms; TE, 30 ms; flip angle, 90°; multiband accelerate factor, 2; voxel size, 2 × 2 × 2 mm; FOV, 224 × 224 mm; 240 volumes for each of three rest scans, 365 and 508 volumes for initial learning and emotional tagging scans separately). Structural images were acquired through three-dimensional sagittal T1-weighted magnetization-prepared rapid gradient echo (MPRAGE) sequence (slices, 192; slice thickness, 1 mm; TR, 2530 ms; TE, 2.98 ms; flip angle, 7°; inversion time, 1100 ms; voxel size, 1 × 1 × 1 mm; FOV, 256 × 256 mm). Field map images were acquired with gre_field_mapping sequence (slices, 64; slice thickness, 2 mm; TR, 635 ms; TE1, 4.92 ms; TE2, 7.38 ms; flip angle, 60°; voxel size, 2 × 2 × 2 mm; FOV, 224 × 224 mm).

### Imaging preprocessing

Brain imaging data was preprocessed using Statistical Parametric Mapping (SPM12; http://www.fil.ion.ucl.ac.uk/spm). The first 4 volumes of functional images were discarded for signal equilibrium. Remaining images were firstly corrected for distortions related to magnetic field inhomogeneity. Subsequently, these functional images were realigned for rigid-body motion correction and corrected for slice acquisition timing. Each participant’s images were then co-registered to their own T1-weighted anatomical image, spatially normalized into a standard stereotactic Montreal Neurological Institute (MNI) space and resampled into 2-mm isotropic voxels. Finally, images were smoothed with a 6-mm FWHM Gaussian kernel.

### Univariate general linear model (GLM) analysis

To assess transient neural activity associated with emotional tagging, we conducted two separate GLMs for initial learning phase and emotional tagging phase. In each GLM, separate regressors of interest were modeled for trials with 2s from onset of each stimulus in aversive condition and neutral condition, and convolved with the canonical hemodynamic response function (HRF) in SPM12. These two conditions were created according to emotional tagging manipulations. During initial learning, trials for face-object pairs subsequently tagged with aversive screams were assigned into aversive condition, and remaining trials with neutral voices were assigned into neutral condition. During emotional tagging, trials tagged with aversive screams were assigned into aversive condition, and trials with neutral voices were assigned into neutral condition. In the GLM of emotional tagging phase, the initial 2-second presentation of each face cue prior to the onset of aversive or neutral voices was also included as a single regressor of no interest. Additionally, each participant’s motion parameters from the realignment procedure were included in each GLM to regress out effects of head movement on brain response. High-pass filtering using a cutoff of 1/128 hz was also included in each GLM to remove high frequency noise and corrections for serial correlations using a first-order autoregressive model (AR(1)).

Contrast parameter estimate images for task-related brain responses in aversive versus neutral conditions from each model were generated at the individual-subject level, and then submitted into a paired t-test for second-level group analysis treating participants as a random factor. Significant clusters were identified from group analyses, initially masked using a gray matter mask, and then determined using a height threshold of *p* < 0.001 and an extent threshold of *p* < 0.05 with family-wise error correction for multiple comparisons based on nonstationary suprathreshold cluster-size distributions computed using Monte Carlo simulations (Nichols & Hayasaka, 2003).

To further investigate specific neural activity associated with the emotion-induced effects on associative memory, we conducted two additional GLMs for initial learning and emotional tagging phases separately. Each GLM included Memory Status (forgotten, remembered with low confidence versus remembered with high confidence) as another variable of interest, together with Emotion (aversive versus neutral). Conditions of Memory Status were created according to associative memory performance. To ensure reliable and strong associative memories, trials later remembered with low confidence regardless of Emotion conditions were included as a single regressor of no interest. Thus five regressors of interest (i.e., later forgotten in aversive condition, later forgotten in neutral condition, later remembered with low confidence regardless of Emotion conditions, later remembered with high confidence in aversive condition, later remembered with high confidence in neutral condition) were modeled in each GLM. In the GLM of emotional tagging phase, face cues with initial 2-second presentation prior to the onset of voices were also included as a regressor of no interest. Relevant parameter contrasts from each model were then submitted to a 2 (Emotion: aversive versus neutral) by 2 (Memory Status: forgotten versus remembered with high confidence) repeated-measures ANOVA for second-level group analysis treating participants as a random factor. All other settings were same as the above GLM analysis.

### Regions of interest (ROIs) definition

To investigate the effects of emotional tagging on memory reactivation, the overall activation mask on whole-brain level was defined using the overlapping area of two group contrast maps of face-object association encoding (initial learning) and face-voice association encoding (emotional tagging) separately relative to fixation, by a very stringent threshold of *p* < 0.0001 with more than 30 voxels. To better characterize hippocampal and face/object-sensitive neocortical reactivation patterns, ROIs in the hippocampus, the face fusiform area (FFA) for face processing (Kanwisher, McDermott, & Chun, 1997) and the lateral occipital cortex (LOC) for object processing (Grill-Spector, Kourtzi, & Kanwisher, 2001) were also separately identified by its corresponding region defined by the Automated Anatomical labelling (AAL) brain atlas and then constrained by the above contrast map. The FFA and LOC were further constrained by overlapping with the mask derived from the Neurosynth platform for large-scale, automated synthesis of fMRI data (http://neurosynth.org/) with “face” and “object” separately as searching term. The two Neurosynth masks were refined by a threshold of *p* < 0.001 with more than 30 voxels. ROI in face/object-sensitive neocortical regions was the combination of FFA and LOC. To further investigate emotion-induced changes in hippocampal functional connectivity during initial learning, emotional tagging, as well as resting state, the above hippocampal ROI was also used as seed in both following task-dependent gPPI and task-free connectivity analyses.

### Multi-voxel pattern similarity analysis

To measure the neural reactivation of initial learning activity for face-object associations cued by the overlapping faces during emotional tagging, we computed multi-voxel pattern similarity of stimulus-evoked activation between initial learning phase and emotional tagging phase for each participant within each ROI. T values of activation maps for aversive and neutral conditions during initial learning and emotional tagging phases were firstly extracted into separate vectors and z-scored. Similarity between vectors of initial learning and emotional tagging phases was computed for each condition using Pearson’s correlation, and then Fisher-transformed. The resultant similarity maps for two conditions were entered into a paired t-test (aversive vs. neutral) for the group-level analysis to determine emotion-induced differences in reactivation between aversive and neutral conditions.

### Whole-brain pattern similarity analysis

A searchlight mapping method was implemented to assess the reactivation of initial learning activity during emotional tagging on the whole-brain level. Similar to above ROI-based analysis, we computed multi-voxel pattern similarity between initial learning and emotional tagging phases for aversive and neutral conditions separately in each searchlight, using a 6-mm spherical ROI centered on each voxel across the whole brain. The resultant Fisher-transformed searchlight maps for two conditions were then entered into a paired-t test (aversive vs. neutral) on the group-level analysis to determine other brain regions involved in emotion-induced increased reactivation. Significant clusters were determined by using the same criterion from the above GLMs.

### Task-dependent functional connectivity analysis

To assess hippocampus-based functional connectivity associated with emotional tagging, we conducted a generalized form of task-dependent psychophysiological interaction (gPPI) analysis during initial learning and emotional tagging phases separately (Friston et al., 1997; S. Qin, Cho, et al., 2014). This analysis examined condition-specific modulation on functional connectivity of a specific ROI (the hippocampal seed here) with the rest of the brain, after removing potentially confounding influences of overall task activation and common driving inputs. The physiological activity of given seed region was computed as the mean time series of all voxels. They were then deconvolved so as to uncover neuronal activity (i.e., physiological variable) and multiplied with the task design vector by contrasting aversive and neutral conditions (i.e., a binary psychological variable) to form a psychophysiological interaction vector. This interaction vector was convolved with a canonical HRF to form the PPI regressor of interest. The psychological variable representing the task conditions (aversive vs. neutral), as well as the mean-corrected time series of the seed ROI, were also included in the GLM to remove overall task-related activation and the effects of common driving inputs on brain connectivity. Contrast images corresponding to PPI effects at the individual-subject level were then submitted to a paired-t test (aversive vs. neutral) on the group-level analysis to determine the effects of emotional tagging on hippocampal functional connectivity with the rest of the brain. Significant clusters were determined using the same criterion from above GLMs. Next, mean t values extracted from the resultant contrast images within each target ROI were also submitted to a 2 (Emotion: aversive vs. neutral) by 2 (Phase: initial learning vs. emotional tagging) repeated-measures ANOVA for the group-level analysis.

To further investigate the effects of subsequent emotional tagging on established associative memory through online hippocampal functional connectivity, we conducted an additional hippocampal-seeded gPPI analysis only during emotional tagging phase by taking Memory Status (i.e., forgotten vs. remembered with high confidence) into account. All procedures were same as the above gPPI analysis, except that we included four PPI regressors of interest (i.e., forgotten in aversive condition, forgotten in neutral condition, remembered with high confidence in aversive condition, remembered with high confidence in neutral condition) into the model. Mean t values extracted from the resultant contrast images within each target ROI were then submitted to a 2 (Emotion: aversive vs. neutral) by 2 (Memory Status: forgotten vs. remembered with high confidence) repeated-measures ANOVA for the group-level analysis.

### Task-free functional connectivity analysis

To assess emotion-related changes in hippocampal functional connectivity during post-learning offline process, we conducted seeded correlational analyses of resting-state fMRI data for three rest scans (i.e., Rest 1, 2 and 3) separately (S. Qin, Young, et al., 2014). Regional time series within the hippocampal seed were extracted from bandpass-filtered images with a temporal filter (0.008 −0.10 Hz), and then submitted into the individual level fixed-effects analyses. A global signal regressor and six motion parameters were included as nuisance covariates to account for physiological and movement-related artifacts. To mitigate potential individual differences in baseline connectivity and examine the specificity of emotion-tagged memory benefit effect on hippocampal connectivity, hippocampal-seeded connectivity map at Rest 1 was subtracted from Rest 2, and connectivity map at Rest 2 was subtracted from Rest 3 for each participant. The resultant connectivity maps were then submitted separately to a second-level multiple regression analysis, with emotion-tagged retroactive memory enhancement (i.e., associative memory performance in the aversive relative to neutral condition) as the covariate of interest. Significant clusters were determined using the same criterion from above GLMs and gPPI analyses.

To further illustrate the relationships between offline hippocampal connectivity and emotion-tagged memory benefit in specific target regions, mean t values were extracted from significant clusters and included in further machine-learning based prediction analysis described below to examine their predictive values to subsequent memory outcomes.

### Prediction analysis

We used a machine-learning prediction approach with balanced fourfold cross-validation (S. Qin, Cho, et al., 2014) to confirm the robustness of the relationship between reactivation (or functional connectivity) and memory performance. This prediction analysis complements conventional regression models which are sensitive to outliers and have no predictive value. Individual’s reactivation (or functional connectivity) index was entered as an independent variable, and their corresponding memory performance was entered as a dependent variable. Data for these two variables were divided into four folds. A linear regression model was built using data from three out of the four folds and used to predict the remaining data in the left-out fold. This procedure was repeated four times to compute a final r_(predicted, observed)_. Such r_(predicted, observed)_, representing the correlation between the observed value of the dependent variable and the predicted value generated by the linear regression model, was estimated as a measure of how well the independent variable predicted the dependent variable. Finally, a nonparametric approach was used to test the statistical significance of the model by generating 1,000 surrogate data sets under the null hypothesis of r_(predicted, observed)_. The statistical significance (*p* value) was determined by measuring the percentage of generated surrogate data.

### Structural equation modeling of brain-behavior relationship

We conducted structural equation modeling (SEM) to further elucidate how emotional tagging affects initial associative memory and its reactivation through functional amygdala-hippocampal-neocortical pathways during online learning using Mplus 7.0 software (https://www.statmodel.com/index.shtml) (Hayes, Preacher, & Myers, 2011). Full mediation models were constructed to investigate how hippocampal-neocortical connectivity mediated the influence of hippocampal-amygdala connectivity on associative memory performance in aversive and neutral conditions separately. We used hippocampal-amygdala connectivity as the predictor, associative memory performance as the outcome, and hippocampal-neocortical connectivity as the mediator. In this model, individual’s hippocampal connectivity with each targeted ROI (i.e., amygdala and face/object-sensitive neocortical regions) was measured with mean t values extracted from the corresponding contrast images of gPPI analysis (later remembered with high confidence vs. forgotten). Individual’s associative memory performance was measured with correctness proportion for face-object associations remembered with high confidence. Moreover, bias-corrected bootstrap was conducted with 1,000 samples to test the mediating effect of hippocampal-neocortical connectivity (Shrout & Bolger, 2002). Both direct and indirect effects of hippocampal-amygdala connectivity on associative memory were estimated, which generated percentile based on confidence intervals (CI).

To investigate how hippocampal-neocortical connectivity mediates the influence of hippocampal-amygdala connectivity on reactivation of initial associative memory in aversive and neutral conditions separately, we constructed additional mediation models with reactivation as the outcome. Individual’s reactivation of each ROI (i.e., hippocampus and face/object-sensitive neocortical regions) was measured with mean similarity index of high-confident remembered associations minus forgotten ones. All other settings were same as the above SEM analysis. All reported *p* values are two-tailed.

### Estimates of effect size

Effect sizes reported for ANOVAs are partial eta squared, referred to in the text as η^2^. For paired t-tests, we calculated Cohen’s *d* using the mean difference score as the numerator and the average standard deviation of both repeated measures as the denominator (Lakens, 2013). This effect size is referred to in the text as d_av_, where ‘av’ refers to the use of average standard deviation in the calculation.

### Data Availability

The data and codes that support the findings of this study are available from the corresponding author upon request.

## Author contributions

Y.Z. and S.Q. conceived the experiment. Y.Z., L.Z. and C.C. conducted the behavioral and fMRI experiments. Y.Z., L.Z. and Y.Z. performed data analysis. Y.Z., G.F. and S.Q. wrote the manuscript.

## Acknowledgements

This work was supported by the National Natural Science Foundation of China (31522028, 81571056), the Open Research Fund of the State Key Laboratory of Cognitive Neuroscience and Learning (CNLZD1503), and the PhD scholarship (<u>201806040186</u>) of the Chinese Scholarship Council.

