## Supplemental information for "Emotional tagging retroactively promotes memory integration through rapid neural reactivation and reorganization"

**Table S1. Acoustic characteristics of the four voice clips**

| | Duration (sec) | Frequency (Hz) | | | | Power (db) | | | |
|---|---|---|---|---|---|---|---|---|---|
| | | median | mean ± s.d. | min | max | median | mean ± s.d. | min | max |
| **Scream1** | 2 | 419.19 | 350.12 ± 131.29 | 99.79 | 477.44 | 78.15 | 77.64 ± 2.75 | 72.91 | 83.52 |
| **Scream2** | 2 | 307.42 | 338.93 ± 82.83 | 198.13 | 514.32 | 78.14 | 74.79 ± 6.51 | 60.07 | 80.76 |
| **“Eh”** | 2 | 205.76 | 205.75 ± 1.42 | 202.92 | 208.14 | 78.15 | 77.40 ± 1.66 | 72.69 | 79.53 |
| **“Ah”** | 2 | 214.52 | 212.14 ± 13.76 | 92.23 | 216.96 | 78.13 | 76.73 ± 9.15 | -5.39 | 79.53 |

Notes: Sec, second; Hz, Hertz; db, decibel.

**Table S2. Brain regions involved in emotion-charged reactivation**

| Brain Regions | Hemisphere | T values | MNI Coordinates | | |
|---|---|---|---|---|---|
| | | | X | Y | Z |
| | | Aversive vs. Neutral | | | |
| Hippocampus | R | 4.88 | 16 | -38 | 8 |
| Parahippocampal gyrus | R | 5.29 | 22 | 6 | -20 |
| Fusiform | L | 3.90 | -36 | -48 | -16 |
| Insula | L | 4.98 | -36 | 4 | -14 |
| | R | 4.70 | 36 | -10 | 16 |
| Inferior frontal gyrus | L | 6.50 | -44 | 20 | -12 |
| | R | 4.93 | 48 | 28 | -6 |
| Middle frontal gyrus | R | 6.35 | 46 | 0 | 58 |
| Superior medial frontal gyrus | L | 5.37 | 0 | 54 | 16 |
| | R | 6.43 | 12 | 28 | 60 |
| Middle temporal gyrus | R | 5.03 | 40 | -60 | 18 |
| Superior temporal gyrus | R | 5.34 | 46 | -34 | 8 |
| Middle occipital lobe | L | 4.55 | -50 | -78 | 8 |
| Precuneus | L | 6.35 | -10 | -60 | 14 |
| Angular gyrus | R | 5.66 | 32 | -62 | 46 |

Notes: Regions were derived from a whole-brain exploratory analysis using searchlight algorithm on neural pattern similarity between initial learning and emotional tagging phases. Only clusters, significant at a height threshold of $p < 0.001$ and an extent threshold of $p < 0.05$ with family-wise error correction for multiple comparisons, are reported with local maximum T statistic in Montreal Neurological Institute (MNI) space. L, left; R, right.

**Table S3. Post-learning hippocampal connectivity changes underlying emotion-tagged retroactive memory enhancement**

| Brain Regions | Hemisphere | T values | MNI Coordinates | | |
|---|---|---|---|---|---|
| | | | X | Y | Z |
| | | | **Rest 2 vs. Rest 1** | | |
| Lateral occipital cortex | L | 4.03 | -42 | -66 | -6 |
| Inferior temporal gyrus | L | 3.72 | -54 | -58 | -10 |
| Lingual gyrus | R | 5.09 | 10 | -88 | -12 |
| | | | **Rest 3 vs. Rest 2** | | |
| Superior frontal gyrus | R | 4.12 | 24 | 52 | 14 |
| Posterior cingulate gyrus | R | 4.35 | 10 | -36 | 40 |
| Inferior parietal lobule | L | 4.22 | -56 | -42 | 40 |
| | R | 4.07 | 52 | -54 | 40 |

Notes: Regions were derived from hippocampal-seeded functional connectivity analyses on three rest scans separately, with emotion-tagged memory enhancement as the covariate of interest. Only clusters, significant at a height threshold of $p < 0.001$ and an extent threshold of $p < 0.05$ with family-wise error correction for multiple comparisons, are reported with local maximum T statistic in Montreal Neurological Institute (MNI) space. L, left; R, right.

**Figure S1**

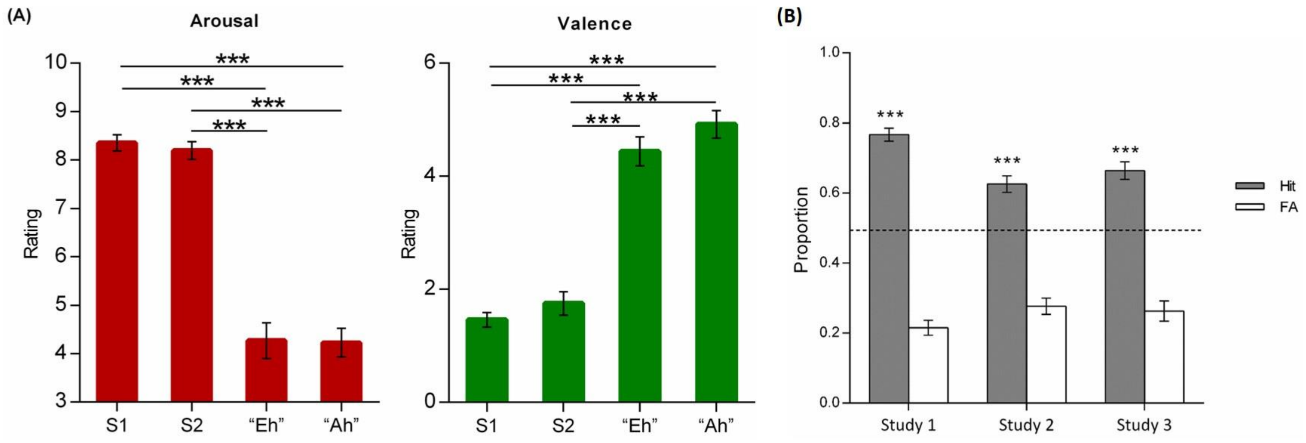

**<u>Figure S1</u>. Emotional ratings on the four voice clips and face item memory performance.**

**(A)** Bar graphs depict averaged arousal (*left*) and valence (*right*) ratings of each voice clip. **(B)** Item memory performances in three studies. About 30 minutes after emotional tagging phase, a self-paced face item memory test was administered prior to face-object associative memory test. A total of 144 faces (72 learnt ones randomly intermixed with 72 novel foils) were presented on the screen one by one. Participants were instructed to make two-alternative forced-choice to indicate whether the presented face was old or new. Bar graphs depict averaged proportions for hit (gray) and false alarm (FA; white) responses in Study 1, 2 and 3 (fMRI). The dashed line represents the chance level (50%). Error bars represent standard error of mean. Notes: S1, Scream 1; S2, Scream 2; ****p* < 0.001; two-tailed tests.

**Figure S2**

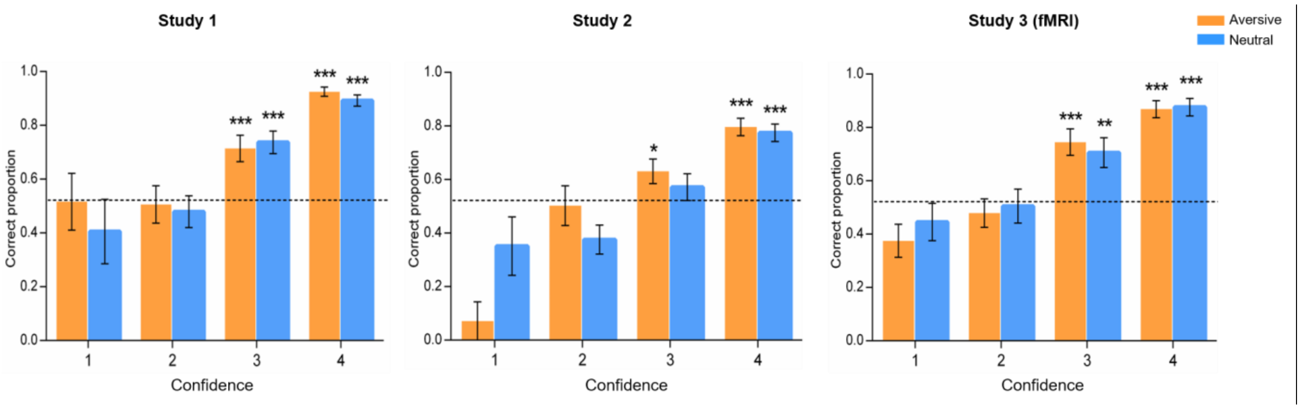

**<u>Figure S2</u>. Associative memory performances on four confidence levels.**

Associative memory performances on each confidence rating, with 1 corresponding to very unconfident and 4 corresponding to very confident, in three studies. Bar graphs depict averaged correctness (correct responses /all responses in each confidence level) for face-object associations in aversive and neutral conditions. In Study 1, separate one-sample t-tests on each confidence level revealed that the proportion of correctly remembered face-object associations with a 4 ($t_{(28)} = 23.00$, $p < 0.001$) or 3 ($t_{(29)} = 5.02$, $p < 0.001$) rating was significantly higher than chance level. However, the proportion of remembered associations with a 2 ($t_{(26)} = -0.29$, $p = 0.771$) or 1 ($t_{(13)} = -0.02$, $p = 0.983$) rating was not reliably different from chance. In Study 2, the proportion of correctly remembered face-object associations with a 4 ($t_{(27)} = 8.59$, $p < 0.001$) or 3 ($t_{(27)} = 2.09$, $p < 0.05$) rating was significantly higher than chance. The proportion of remembered associations with a 2 ($t_{(24)} = -2.30$, $p = 0.031$) or 1 ($t_{(9)} = -2.68$, $p = 0.025$) rating was even lower than chance. In Study 3, the proportion of correctly remembered face-object associations with a 4 ($t_{(26)} = 12.48$, $p < 0.001$) or 3 ($t_{(27)} = 3.83$, $p < 0.001$) rating was significantly higher than chance. The proportion of remembered associations with a 2 ($t_{(25)} = -0.52$, $p = 0.607$) or 1 ($t_{(17)} = -2.53$, $p = 0.022$) rating was not different from or even lower than chance. The dashed lines represent the chance level (52%). Error bars represent standard error of mean. Notes: $^{**}p < 0.01$, $^{***}p < 0.001$; two-tailed one-sample t-tests.

**Figure S3**

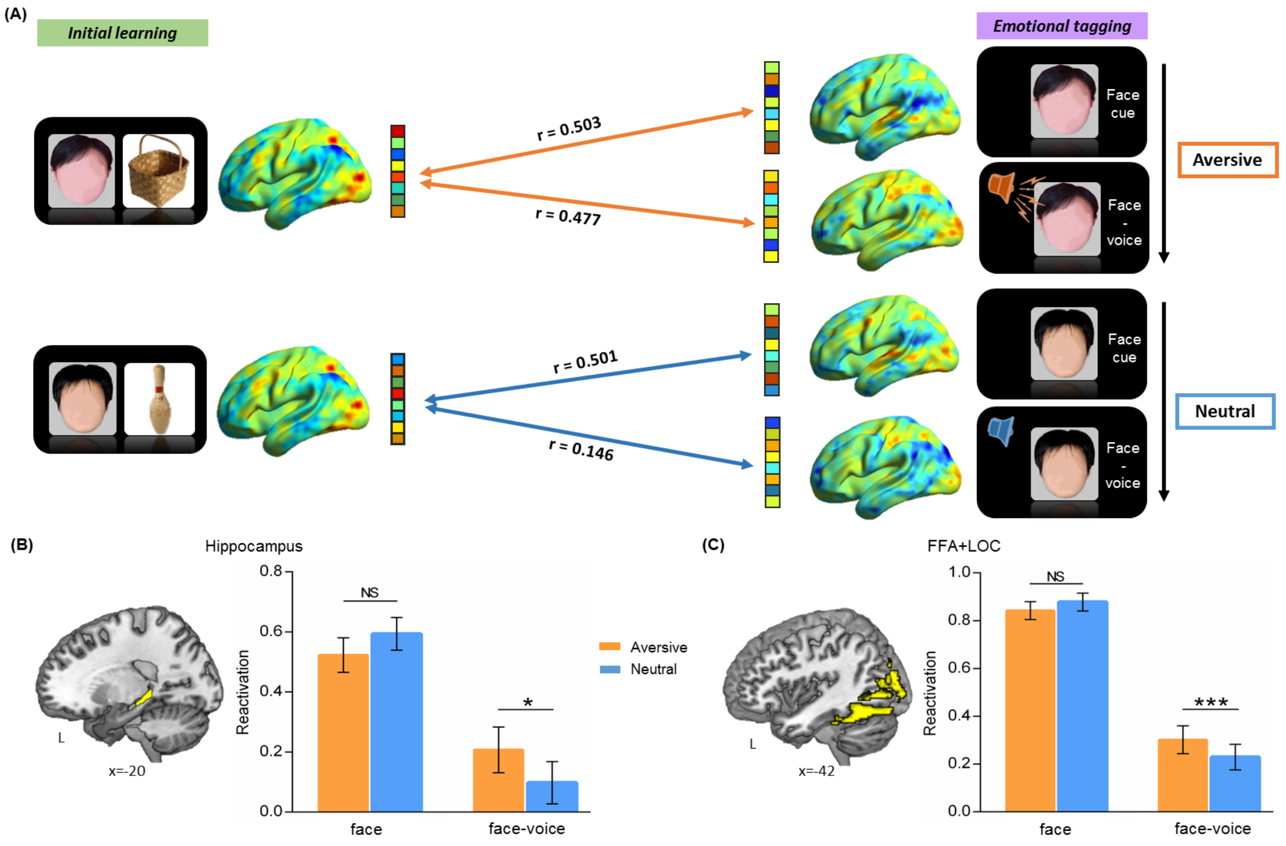

**Figure S3. Neural reactivation of initial learning activity during face cue only and face-voice association separately in emotional tagging phase.**

**(A)** An illustration of reactivation analysis by computing similarity of stimulus-evoked multi-voxel activity patterns between initial learning and emotional tagging phases. Example data was from one subject. During initial learning (*left*), sagittal views of activation maps for aversive and neutral conditions were shown. During emotional tagging (*right*), sagittal views of activation maps for two conditions during the presentation of face cue only (*upper*) and the following presentation of face-voice association (*lower*) were shown separately. Faces in the figure are completely obscured for copyright reasons. **(B, C)** Bar graphs depict averaged reactivation of the hippocampus and face/object-sensitive neocortical regions (e.g. FFA and LOC) in aversive and neutral conditions during face cue only and face-voice association separately. There was no significant difference in reactivation between aversive and neutral conditions during face cue only (both $t_{(27)} < -1.50$, $p > 0.05$), but significantly higher reactivation in both the hippocampus ($t_{(27)} = 2.28$, $p = 0.03$, $d_{av} = 0.28$) and neocortical regions ($t_{(27)} = 4.41$, $p < 0.001$, $d_{av} = 0.25$) during face-object association in aversive (vs. neutral) condition. Error bars represent standard error of mean. Notes: NS, non-significant; *$p < 0.05$; ***$p < 0.001$; two-tailed t-tests; FFA, fusiform face area; LOC, lateral occipital cortex; L, left.

**Figure S4**

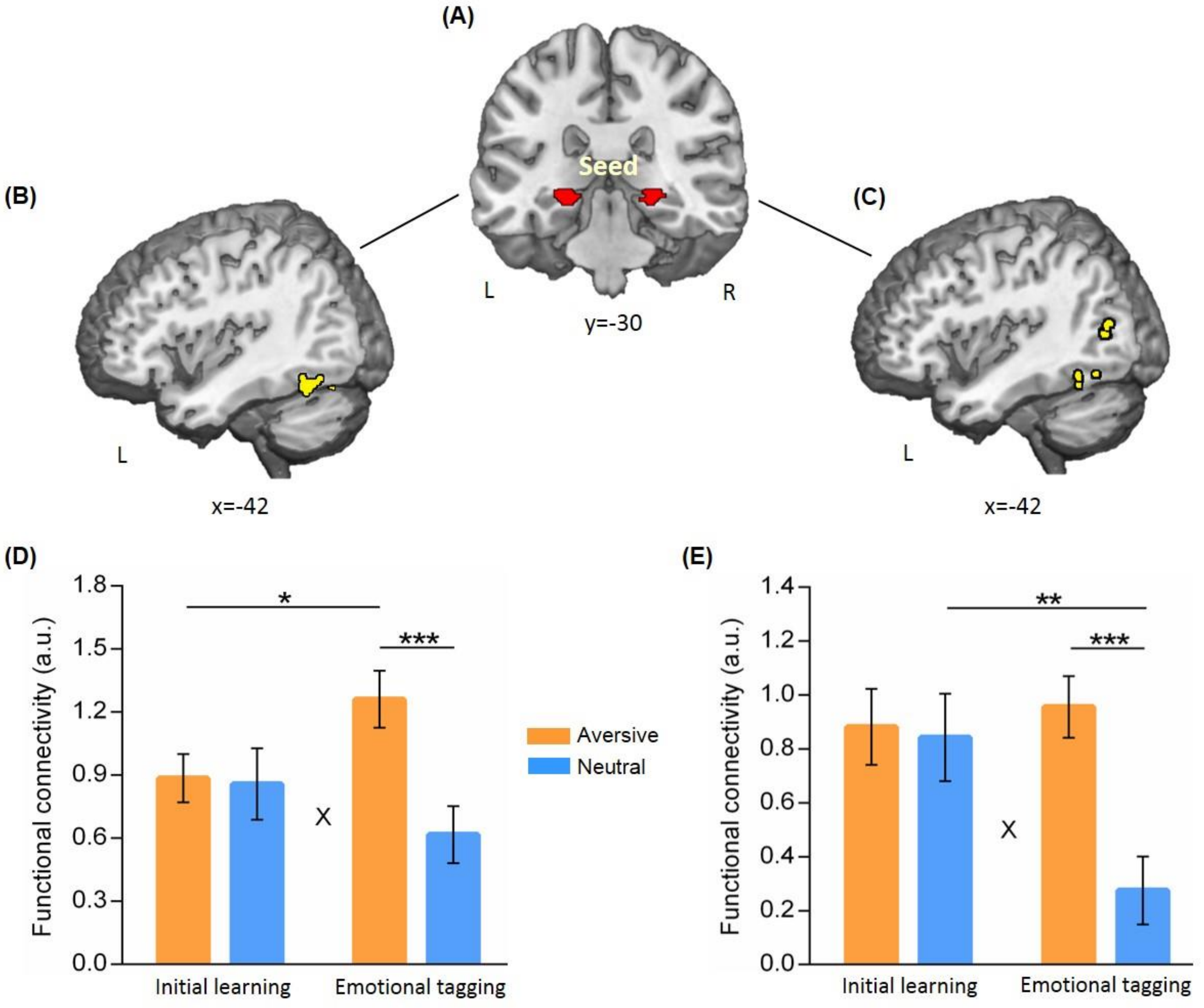

**<u>Figure S4</u>. Hippocampal connectivity with the FFA and the LOC during initial learning and emotional tagging.**

**(A)** Bilateral hippocampal seed used in task-dependent gPPI functional connectivity analyses. **(B)** Significant clusters in the left FFA, and **(C)** significant clusters in the left LOC, showing their greater functional connectivity with the hippocampus in aversive (vs. neutral) condition during emotional tagging phase. **(D, E)** Bar graphs depict averaged hippocampal connectivity with the FFA and the LOC in aversive and neutral conditions during initial learning and emotional learning phases separately. Error bars represent standard error of mean. Notes: *$p < 0.05$; **$p < 0.01$; ***$p < 0.001$; two-tailed tests; L, left; R, right.

**Figure S5**

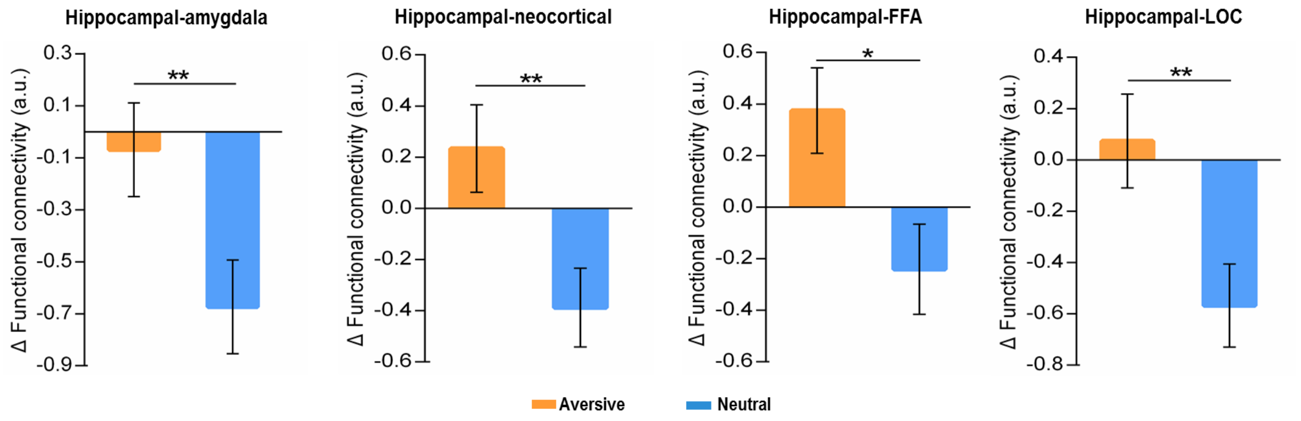

**<u>Figure S5</u>. Hippocampal connectivity during emotional tagging relative to initial learning in aversive and neutral conditions.**

Bar graphs depict averaged hippocampal connectivity with the amygdala, neocortical regions (i.e., FFA and LOC), the FFA and the LOC during emotional tagging phase relative to initial learning phase in two conditions. Notes: *$p < 0.05$; **$p < 0.01$; two-tailed tests.

**Figure S6**

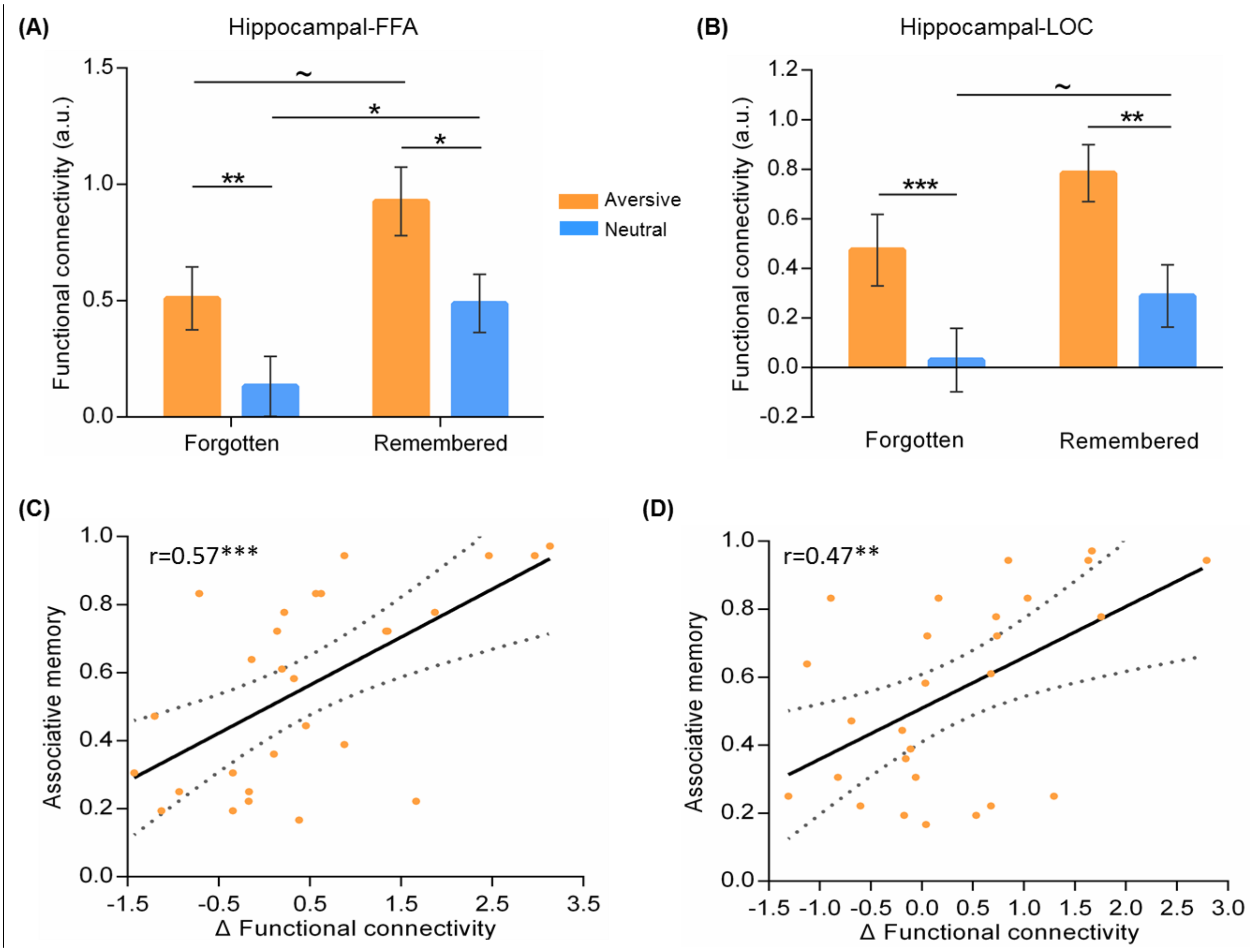

**Figure S6. Hippocampal connectivity with the FFA and the LOC in relation to associative memory performance during emotional tagging.**

**(A, B)** Bar graphs depict averaged hippocampal connectivity with the FFA and the LOC for face-object associations remembered with high confidence and forgotten in aversive and neutral conditions separately. **(C, D)** Scatter plots depict positive correlations of hippocampal-FFA and hippocampal-LOC connectivity (remembered with high confidence vs. forgotten) with associative memory performance in aversive condition. Dashed lines indicate 95% confidence intervals, and solid lines indicate the best linear fit. Error bars represent standard error of mean. Notes: ~ $p < 0.08$; *$p < 0.05$; **$p < 0.01$; ***$p < 0.001$; two-tailed tests.

**Figure S7**

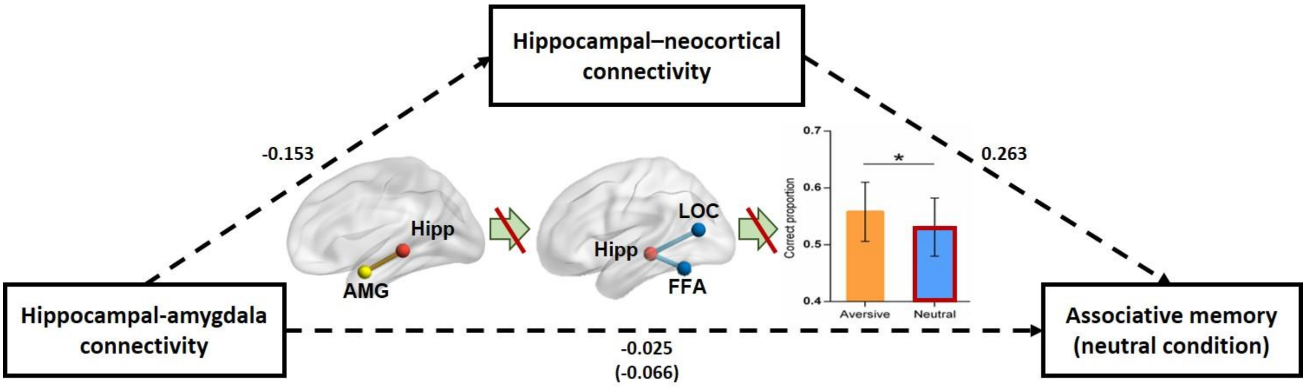

**<u>Figure S7</u>. The mediating effect of online task-dependent hippocampal-neocortical connectivity on the association between hippocampal-amygdala connectivity and associative memory in neutral condition.**

The mediating effect was non-significant (Indirect Est = -0.04, $p$ = 0.602, 95% CI = [-0.19, 0.11]). Paths are marked with standardized coefficients. Dashed lines indicate non-significant paths. The coefficient in bracket shows the correlation before hippocampal-neocortical connectivity was included into this model. Notes: *$p$ < 0.05; Hipp, hippocampus; AMG, amygdala.

**Figure S8**

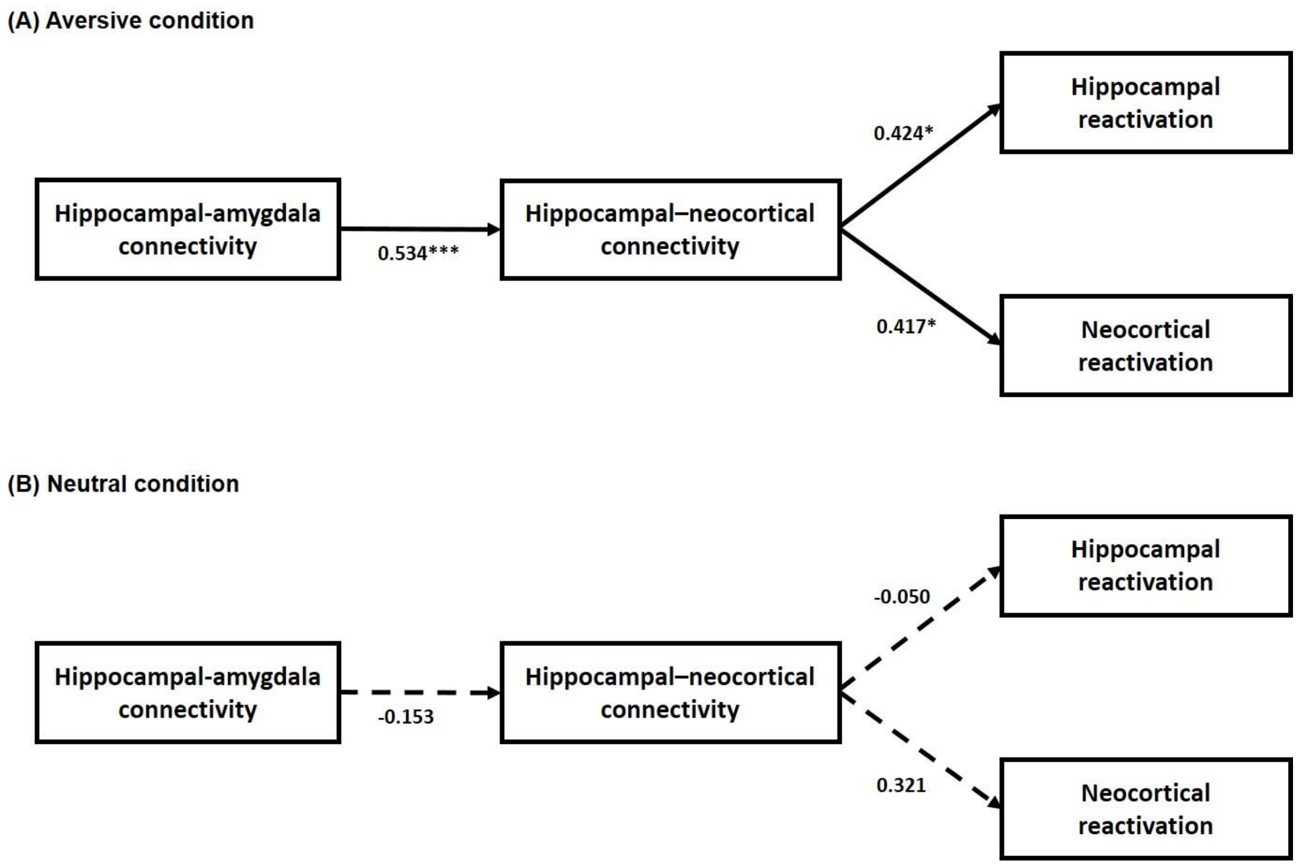

**Figure S8. The mediating effect of online task-dependent hippocampal-neocortical connectivity on the association of hippocampal-amygdala connectivity with hippocampal and neocortical reactivation.**

**(A)** Mediation model in aversive condition. This model resulted in good model fit (Chi$^2$ = 1.455, $p$ > 0.050; RMSEA = 0.000; SRMR = 0.050; CFI = 1.00). The mediating effects of hippocampal-neocortical connectivity, with hippocampal reactivation (Indirect Est = 0.226, $p$ = 0.042, 95% CI = [0.008, 0.444]) and neocortical reactivation (Indirect Est = 0.222, $p$ = 0.040, 95% CI = [0.010, 0.435]) as the outcome separately, were both significant. **(B)** Mediation model in neutral condition. The model fitting coefficients are: Chi$^2$ = 2.560, $p$ > 0.050; RMSEA = 0.102; SRMR = 0.070; CFI = 0.641. The mediating effects, with hippocampal reactivation (Indirect Est = 0.008, $p$ = 0.800, 95% CI = [-0.051, 0.057]) and neocortical reactivation (Indirect Est = -0.049, $p$ = 0.547, 95% CI = [-0.209, 0.085]) as the outcome separately, were both non-significant. Paths are marked with standardized coefficients. Solid lines indicate significant paths, and dashed lines indicate non-significant paths. Notes: *$p$ < 0.05; ***$p$ < 0.001; two-tailed tests.
